# The stability and ABA import activity of NRT1.2 in *Arabidopsis* are regulated by CEPR2 via phosphorylation modification

**DOI:** 10.1101/2020.02.18.953711

**Authors:** Lei Zhang, Zipeng Yu, Yang Xu, Miao Yu, Yue Ren, Shizhong Zhang, Guodong Yang, Jinguang Huang, Kang Yan, Chengchao Zheng, Changai Wu

## Abstract

Abscisic acid (ABA) transport plays important role in systematic plant responses to environmental factors. Here, we showed that C-terminally encoded peptide receptor 2 (CEPR2) directly interacted with the ABA transporter NRT1.2. Using transgenic seedlings, we demonstrated that NRT1.2 positively regulated the ABA response, and that CEPR2 acted on NRT1.2 epistatically and negatively. Under normal conditions, CEPR2 phosphorylated NRT1.2 at least at serine 292 to promote the degradation of NRT1.2. However, ABA and serine 292 nonphospharylation strongly inhibited the degradation of NRT1.2, indicating that ABA-inhibited the phosphorylation of NRT1.2. Transport assays in yeast and *Xenopus* oocytes showed that nonphosphorylated NRT1.2 had high levels of ABA-import activity, but phosphorylated NRT1.2 did not import ABA. Analyses of complement *nrt1.2* mutants by mimicking nonphospharylated and phospharylated NRT1.2 confirmed that nonphospharylated NRT1.2^S292A^ had high stability and ABA import activity *in planta*. Further experiments indicated that NRT1.2 was degraded via the 26S proteasome and vacuolar degradation pathways. UBC32, UBC33, and UBC34 interacted with, and mediated the ubiquitination of NRT1.2. UBC32, UBC33, and UBC34 acted on NRT1.2 epistatically and negatively *in planta*. Thus, our results suggested the existence of novel plant mechanisms regulating NRT1.2 stability and ABA import activity in response to environmental conditions.

## Introduction

Plants must continuously adapt to adverse environmental conditions, including temperature, salinity, and drought (Li et al., 2019; Soma et al., 2017; Teardo et al., 2019; Xu et al., 2018; Yang et al., 2019; Zhang et al., 2018). Upon sensing environmental stressors, plants stop growing and trigger protective responses (Wang et al., 2018). Once environmental stress subside, plants quickly deactivate protective responses and reinitiate growth (Wang et al., 2018). Thus a rapid response to an environmental stress is critical for plant growth and survival. However, the mechanisms underlying these rapid responses to adverse conditions are not fully understood.

Abscisic acid (ABA), a stress hormone, plays critical roles in the regulation of both plant growth and numerous stress responses (Cutler et al., 2010; Raghavendra et al., 2010). In response to environmental stress, ABA accumulates rapidly and is transported from the site of synthesis to the site of action, where it is recognized by ABA receptors in the pyrabactin resistance 1 family (PYR1); these proteins are also referred to as PYR1-like (PYL) (Boursiac et al., 2013). ABA is predominantly produced in vascular tissues in response to drought stress (Boursiac et al., 2013). Therefore, ABA signaling in plants appears to rely on the export of ABA from ABA-biosynthesis cells as well as the import of ABA into target cells (Kuromori et al., 2018). Several ABA transporters have been identified. Two *Arabidopsis* AT-binding cassette (ABC) transporters, AtABCG25 and AtABCG40, have been identified as ABA transporters (Kang et al., 2015; Kuromori et al., 2010). Using mutant plants where the expression of these transporters was inhibited, AtABCG25 and AtABCG40 were shown to be associated with ABA sensitivity and stomatal closure (Kang et al., 2015; Kuromori et al., 2010). Additional biochemical characterization studies, using heterologous expression systems, have shown that AtABCG25, AtABCG30, AtABCG31, and AtABCG40 exhibit ABA export or import activity. Furthermore, a member of the multidrug and toxin efflux (MATE) transporter family, AtDTX50, has also been identified as an ABA exporter (Zhang et al., 2014). Finally, functional screening using a yeast system identified an Arabidopsis NITRATE TRANSPORTER 1/PEPTIDE TRANSPORTER FAMILY (NPF) member, NPF4.6 (originally named AIT1), that mediated the cellular uptake of ABA (Kanno et al., 2012). Because NPF4.6 was previously characterized as a low-affinity nitrate transporter NRT1.2 (Huang et al., 1999), we refer to this protein as NRT1.2 herein. However, the mechanisms underlying ABA transporter function, particularly with respect to posttranslational regulation, have not been explored.

Previously, we reported that CEPR2, a typical leucine-rich repeat receptor-like kinase (LRR-RLK), controls the ABA response in *Arabidopsis* by regulating the phosphorylation and degradation of ABA-receptor PYLs (Yu et al., 2019). These results demonstrated, for the first time, that plasma-membrane-localized LRR-RLK regulated ABA receptors. However, a thorough understanding of the mechanisms underlying CEPR2-mediated ABA response in *Arabidopsis* is lacking. Therefore, in this study, we aimed to determine whether CEPR2 regulates ABA transporters. Here, we show that the CEPR2-mediated ABA response was activated by the ABA-inhibited phosphorylation of NRT1.2 at the residue serine 292 (Ser292). Therefore, our results identified a novel target of CEPR2 and characterized the regulatory mechanisms underlying ABA transporter activity as well as NRT1.2 stability.

## Results

### The loop region of NRT1.2 interacts with the kinase domain of CEPR2

To identify new targets of CEPR2 in response to ABA, we compared the phosphoproteomes (Supplementary Fig.1A) of wild-type (WT) *Arabidopsis* seedlings to those of CEPR2-OE-9 seedlings; CEPR2-OE-9 is a previously-developed *Arabidopsis* line overexpressing CEPR2 (Yu et al., 2019). Across both groups of seedlings, 169 differentially regulated phosphopeptides were found (where P ≤ 0.05 and fold-change ≥ 1.20 or ≤ 0.8). Of these 97 (belonging to 78 proteins) were upregulated and 72 (belonging to 68 proteins) were downregulated (Supplementary Excel 1). To identify the directly-phosphorylated target proteins of CEPR2, we tested whether the 78 phosphorylation-upregulated proteins (Supplementary Table 1) interacted with CEPR2 using yeast two hybrid (Y2H) experiments with the mating-based split ubiquitin system (MbSUS) (Supplementary Fig. 1). Ten of the proteins interacted with CEPR2. Of these, NRT1.2 (protein number 46) has previously been identified as an ABA importer (Huang et al., 1999; Kanno et al., 2012). Thus, NRT1.2 was chosen for further analysis.

To confirm the interaction between NRT1.2 and CEPR2, firefly luciferase complementation imaging (LCI) assays were performed in *Nicotiana benthamiana* leaves. The results were consistent with the MbSUS results which showed that NRT1.2 interacted with CEPR2 (Fig. 1A). Next, the split YFP combinations (CEPR2-YFPn and NRT1.2-YFPc) were transiently co-expressed in *N.benthamiana* leaves. Bimolecular fluorescence complementation (BiFC) assays, in conjunction with N-(3-triethylammomiumpropyl) 4-(p-diethylaminophenylhexa-trienyl) (FM4-64) staining, indicated that NRT1.2 and CEPR2 interacted in the plasma membrane (Fig. 1B). Differential phosphoproteome analysis showed that the phosphorylation sites of NRT1.2 were the residues serine 277 and 292 (Ser^277^ and Ser^292^) (Supplementary Excel 1). These two sites were located in the cytosol loop region of NRT1.2 between two of the 12 transmembrane regions as shown in Fig. 1C. To further test the interaction between NRT1.2 and CEPR2, we performed pulldown assays using the cytosol loop region of NRT1.2 (NRT1.2^loop^) and CEPR2 series constructs developed previously (Yu et al., 2019). The results indicated that the NRT1.2^loop^ interacted with the kinase domain of CEPR2 (CEPR2^KD^) (Fig. 1D). Next, NRT1.2^loop^-GST and CEPR2^KD^-His were expressed in *Escherichia coli* and purified. The results of the pulldown assay also showed that NRT1.2^loop^ interacted with CEPR2^KD^ (Fig. 1E). Finally, the genes encoding ABA transporters were not significantly differentially expressed among CEPR2-OE-9, WT, and *cepr2-1* seedlings grown on 1/2 MS with or without 1 μM ABA media. The *cepr2-1* seedlings were developed previously, see Yu et al., 2019 (Supplementary Fig. 2). These genes also did not interact with CEPR2 (Fig. 1F). Thus, four independent approaches, MbSUS, BiFC, LCI, and pulldown, all demonstrated an interaction between the NRT1.2 ^loop^ region and the CEPR2 kinase domain *in vitro* and in plant cells.

**Figure 1.**
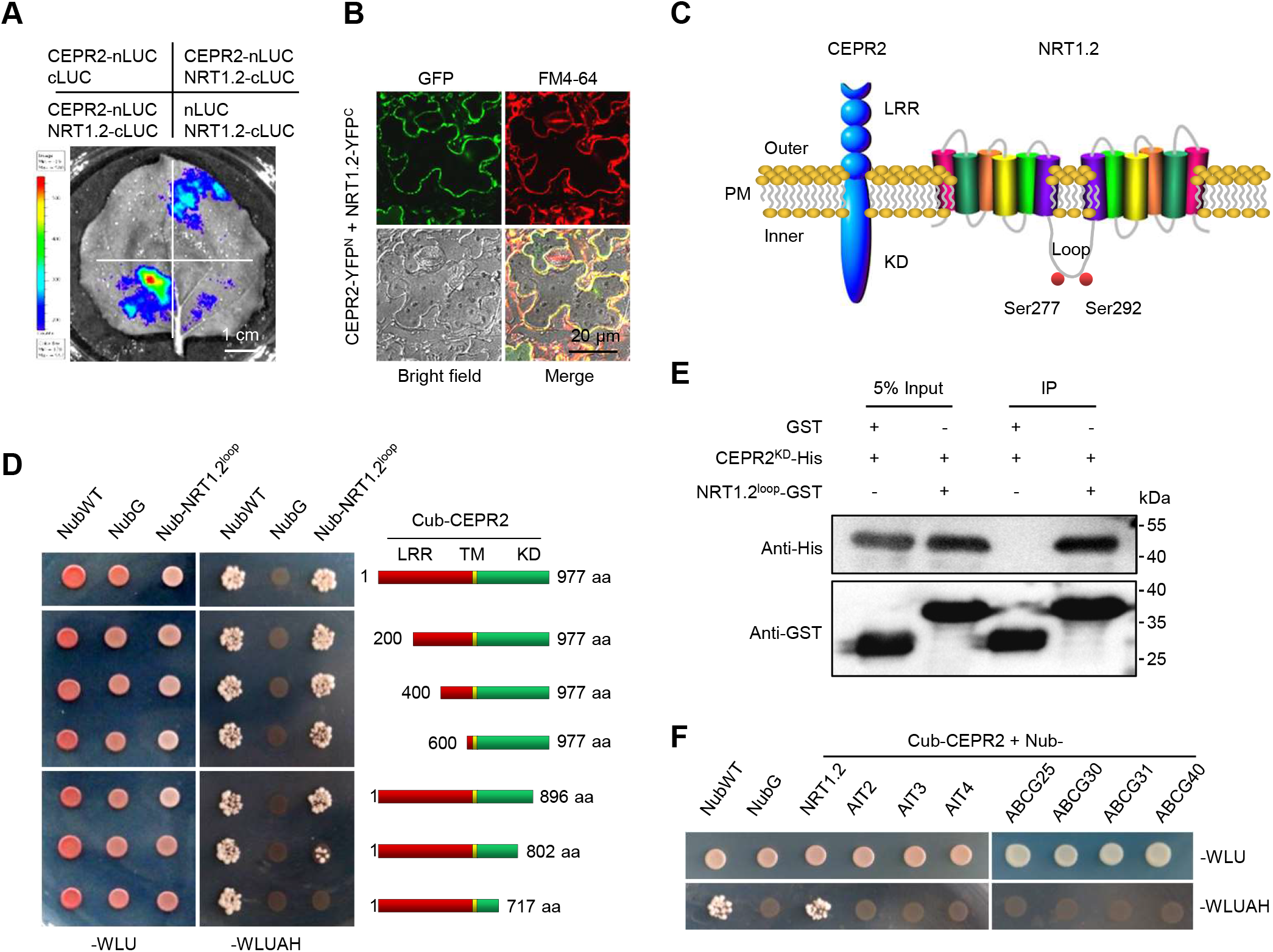
CEPR2 interacts with the ABA importer NRT1.2. **(A)** Firefly luciferase complementation imaging assay, showing the interactions between CEPR2 and NRT1.2. A key to the combination shown in each quadrant is given above the image. **(B)** Bimolecular fluorescence complementation assay, showing the interaction between CEPR2 and NRT1.2 in Nicotiana benthamiana. FM4-64, plasma membrane dye N-(3-triethylammomiumpropyl) 4-(p-diethylaminophenylhexa-trienyl). **(C)** The protein structure of CEPR2 and NRT1.2. The phosphorylation sites S277 and S292 identified by phosphorylation mass spectrometry are labeled in the NRT1.2 protein. LRR, leucine-rich repeat receptor-like domain; KD, kinase domain; Loop, loop domain of NRT1.2; PM, plasma membrane. **(D)** Mating-based split ubiquitin system assay (MbSUS) assay showing the functional domains of CEPR2 that interact with the loop region of NRT1.2 (NRT1.2^loop^, 232–346 aa). **(E)** Pulldown assays using purified GST, NRT1.2loop-GST, and CEPR2KD-His expressed in *E.coli*. **(F)** MbSUS assay showing the interaction between CEPR2 and eight of characterized ABA importers, including NRT1.2, ABA importer 2 (AIT2), AIT3, AIT4, ABC transporter G25 (ABCG25), ABCG30, ABCG31, and ABCG40.

### NRT1.2 positively regulates the ABA-mediated inhibition of seed germination and primary root growth

To assess the role of NRT1.2 in the ABA response, two independent Arabidopsis lines overexpressing *NRT1.2* (NRT1.2-OE-1 and NRT1.2-OE-2) (Fig. 2A) and two T-DNA insertion mutants of NRT1.2 (*nrt1.2-1* and *nrt1.2-2*) were used (Fig. 2B). Reverse transcription-polymerase chain reactions (RT-PCRs) and western blots demonstrated that NRT1.2 was overexpressed in the transgenic NRT1.2-OE-1 and NRT1.2-OE-2 plants. Although the expression levels of NRT1.2 in the *nrt1.2-2* seedlings were similar to those in the WT seedlings, NRT1.2 protein levels in the two mutant seedlings were substantially lower than those in the WT seedlings (Fig. 2C). Under normal conditions, the NRT1.2-OE and *nrt1.2* plants exhibited similar primary root growth and seed germination rates to the WT (Fig. 2D, E and F). When treated with ABA, however, the NRT1.2-OE plants had shorter primary roots and lower seed germination rates than the WT plants. In contrast, the *nrt1.2* mutant plants developed more quickly, and had longer primary roots (Fig. 2D-F). These results suggested that NRT1.2 positively regulates the ABA response.

**Figure 2.**
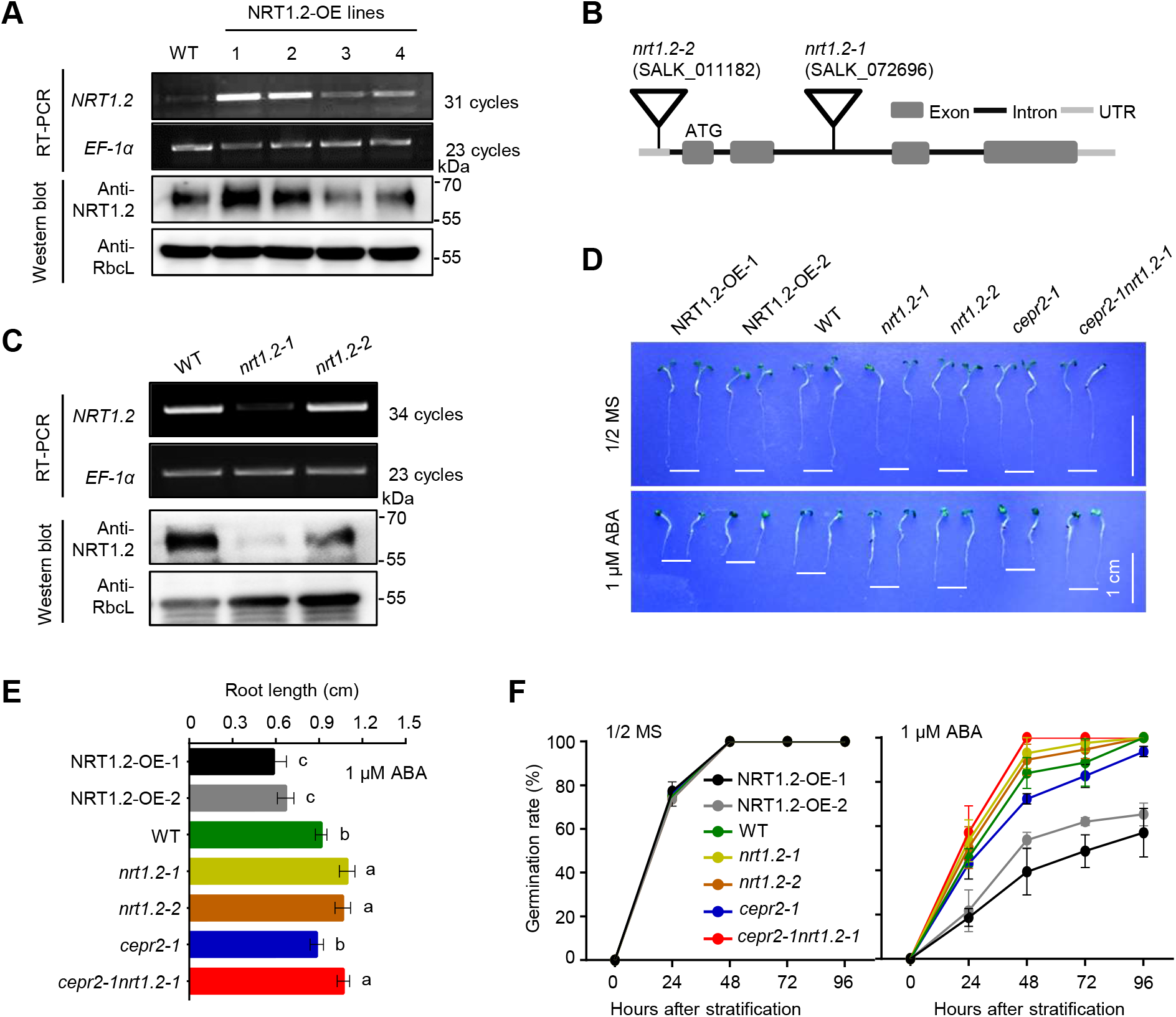
CEPR2 acts epistatically on NRT1.2 to affect ABA signaling. **(A)** NRT1.2 transcript levels and protein levels in WT and transgenic Arabidopsis lines overexpressing NRT1.2 detected using RT-PCRs and western blot. *EF-1α*. was used as the internal control for RT-PCR, and Rubisco large subunit (RbcL) was used as a loading control for the western blot. **(B)** Schematic illustration of the T-DNA insertion sites in the *nrt1.2-1* and *nrt1.2-2* mutants. **(C)** NRT1.2 transcript levels and protein levels in WT, *nrt1.2-1*, and *nrt1.2-2* seedlings. **(D)** Phenotypes of the WT, NRT1.2-OE, *nrt1.2-1, nrt1.2-2, cepr2-1*, and *cepr2-1nrt1.2-1* seedlings grown for seven days on 1/2 MS with or without 1 μM ABA. **(E)** Lengths of the primary roots of the seedlings shown in (D). Error bars indicate SEM (N = 3). Bars labeled with different lowercase letters are significantly different from one another (P < 0.05; one way ANOVA). **(F)** The germination rates of the seedlings shown in (D). Error bars indicate SEM (N = 3).

### CEPR2 phosphorylates NRT1.2 at Ser292, playing a vital role in maintaining NRT1.2 stability

To address the effects of CEPR2 on NRT1.2, genetic phenotype analysis was performed. The results showed that the root length and seed germination rate*s* of the *cepr2-1nrt1.2-1* double mutant were similar to those of the *nrt1.2-1* single mutant, but longer than those of the *cepr2-1* mutant (Fig. 2D-F). This genetic evidence, in conjunction with the results showing that CEPR2 and NRT1.2 interacted, suggested that CEPR2 acts epistatically and negatively on NRT1.2, possibly by regulating the stability or import activity of this protein. To test this possibility, cell-free degradation assays were performed using NRT1.2^loop^-GST with protein extracts from CEPR2-OE-9, WT, and *cepr2-1* seedlings. Under normal conditions, about 53%, 68%, and 12% of the NRT1.2^loop^-GST remained after a 60 min incubation with protein extracts from WT, *cepr2-1* and CEPR2-OE-9 seedlings, respectively (Fig. 3A, B). This indicated that CEPR2 promotes the degradation of NRT1.2 under normal conditions. In the presence of ABA, about 63%, 80%, and 48% of the NRT1.2^loop^-GST remained after a 60 min incubation with protein extracts from WT, *cepr2-1* and CEPR2-OE-9 seedlings, respectively (Fig. 3A, B). These results suggested that CEPR2 decreases the stability of NRT1.2.

**Figure 3.**
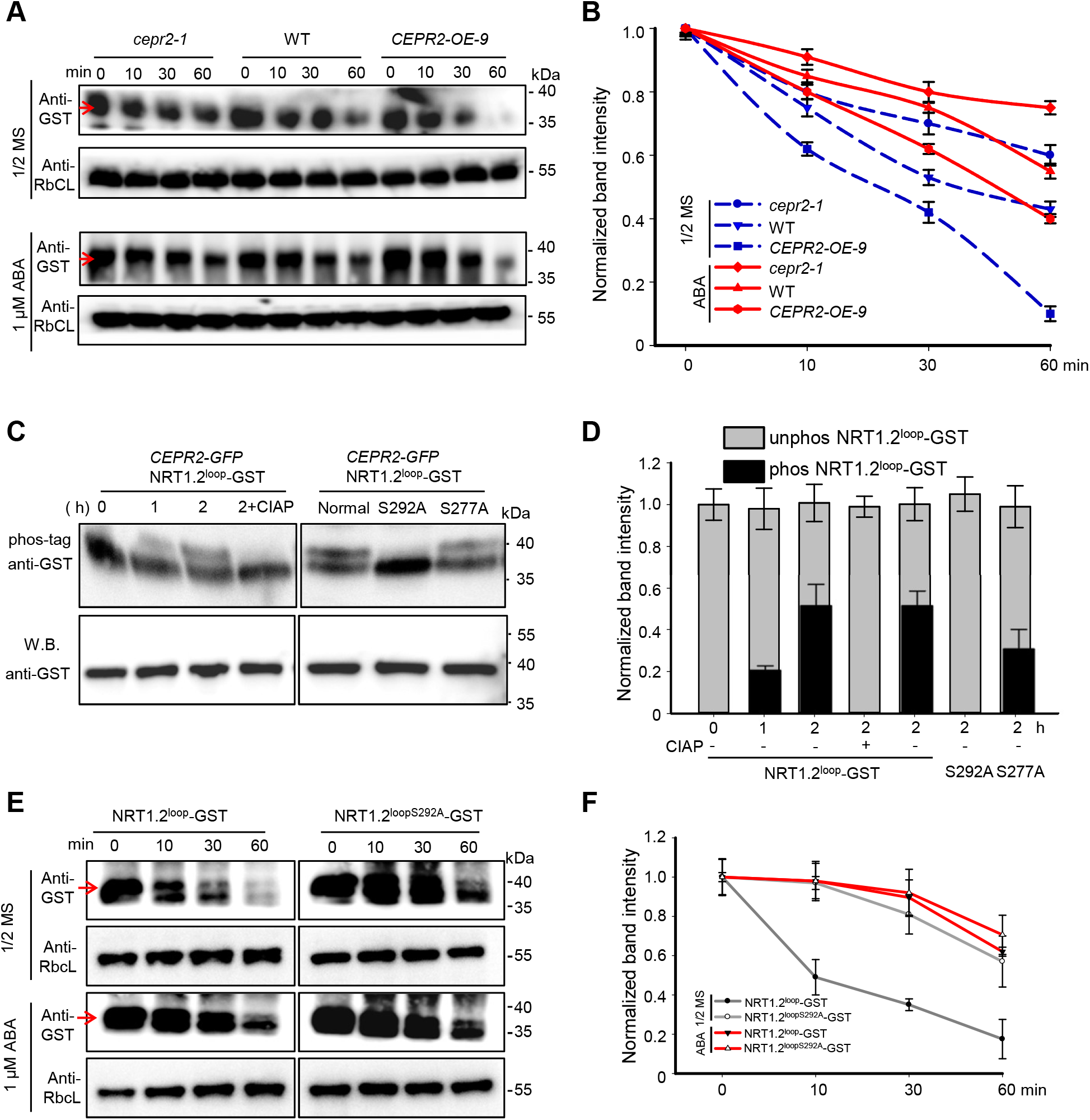
CEPR2 reduces NRT1.2 stability depending on the phosphorylation of Serine 292 (Ser292). **(A)** The cell-free degradation of NRT1.2-GST in soluble protein extracts of CEPR2-OE-9, WT, and cepr2-1 seedlings treated with or without 1 μM ABA. NRT1.2loop-GST protein levels were quantified using anti-GST antibodies (indicated by red arrows). **(B)** Normalized plots of the NRT1.2 degradation shown in (A), based on the band intensities shown in (A); band intensities were quantified using ImageJ as described in methods. Error bars indicate SEM (N = 3). **(C)** In vitro kinase assays showing the phosphorylation effects of CEPR2 on the loop regions of NRT1.2 (NRT1.2^loop^-GST, NRT1.2^loop^S292A-GST and NRT1.2^loop^S277A-GST). NRT1.2 phosphorylation status was detected based on the phos-tag. Calf intestinal alkaline phosphatase (CIAP) was used to remove phosphate group(s) of the proteins via dephosphorylation. **(D)** Normalized plot showing the relative contents of phosphorylated and nonphosphorylated NRT1.2, based on the band intensities shown in (C); band intensities were quantified using ImageJ as described in methods. Error bars indicate SEM (N = 3). Bars labeled with different lowercase letters are significantly different from one another (P < 0.05; one way ANOVA). **(E)** Cell-free degradation assays showing the effects of ABA on the degradation of NRT1.2^loop^ and NRT1.2^loopS292A^ in the soluble protein extracts from CEPR2-OE-9 seedlings. NRT1.2^loop^-GST and NRT1.2^loopS292A^-GST were quantified using western blots with the anti-GST antibody. RbcL was used as the loading control. **(F)** Normalized plot of the NRT1.2 contents based on the band intensities shown in (E); band intensities were quantified using ImageJ as described in methods. Error bars indicate SEM (N = 3).

Phosphorylation modification regulates protein stability, as well as protein activity, interaction, and subcellular localization (Chen et al., 2018). Therefore, the effects of CEPR2 on NRT1.2 stability might be regulated by the phosphorylation states at Ser^277^ and/or Ser^292^. First, we confirmed that CEPR2 interacted with forms of NRT1.2 where Ser^277^ and/or Ser^292^ were mutated to similarly-sized amino acids: non-phosphorylatable, neutral alanine (A) and sustainably phosphorylatable, polar aspartate (D); these forms were referred as NRT1.2^loopS277A^, NRT1.2^loopS292A^, NRT1.2^loopS277D^, NRT1.2^loopS292D^, NRT1.2^loopS277AS292A^, and NRT1.2^loopS277DS292D^ (Supplementary Fig. 3 and 4A). These interactions indicated that mutations in NRT1.2 at Ser^277^ and/or Ser^292^ did not affect the interaction with CEPR2. Therefore, we purified the CEPR2-GFP fusion proteins from CEPR2-GFP transgenic seedlings previously obtained (Yu et al., 2019). Optimized NRT1.2^loop^ proteins from *E.coli* were used to perform *in vitro* kinase assays. When CEPR2-GFP was incubated with NRT1.2^loop^-GST in kinase buffer, the phosphorylation of NRT1.2^loop^-GST increased (Fig. 3C), whereas calf intestinal alkaline phosphatase (CIAP) treatment decreased the phosphorylation of NRT1.2^loop^-GST. This suggested that NRT1.2loop-GST was phosphorylated by CEPR2.

We then incubated CEPR2-GFP with NRT1.2 mutated at Ser^277^ and/or Ser^292^. Subsequent *in vitro* kinase assays showed that, after 2 h, the increased phosphorylation of NRT1.2^loopS277A^-GST was similar to that of NRT1.2^loop^-GST. However, no phosphorylation was detected in NRT1.2^loopS292A^-GST (Fig. 3C and D). These data indicated that Ser^292^, but not Ser^277^, was successfully phosphorylated by CEPR2.

Next, we compared the degradation of NRT1.2^loopS292A^ to that of NRT1.2^loop^. We found that degradation of NRT1.2^loopS292A^ was significantly reduced compared with that of NRT1.2^loop^ irrespective of ABA treatment (Fig. 3E and F). That is, the level of NRT1.2^loopS292A^ in the absence of ABA was similar to the level of NRT1.2^loop^ in the presence of ABA. These results suggested that residue Ser^292^ of NRT1.2 plays a vital role in the phosphorylation-dependent degradation of NRT1.2.

### NRT1.2 phosphorylation eliminates NRT1.2-mediated ABA import

We further investigated the effect of NRT1.2 nonphosphorylation on ABA import activity. Following a previously described Y2H method (Kanno et al., 2012), yeast cells were co-transformed with AD-ABI1 (ABI1 fused to the GAL4 activation domain) and BD-PYR1 (PYR1 fused to the GAL4 DNA-binding domain), and used as background. Compared with pYES2-transformed yeast cells, pYES2-NRT1.2-transformed yeast cells grew better in the presence of 1 μM ABA (Supplementary Fig. 4B). However, no obvious growth differences were observed between pYES2 and pYES2-NRT1.2 yeast cells in the presence of 4 μM ABA. Thus, it was necessary to determine a suitable ABA concentration for subsequent experiments.

None of the transformed yeast cells grew in - WLUAH media. However, in the presence of 1 μM ABA, the yeast cells transformed with nonphosphomimic forms of NRT1.2, such as pYES2-NRT1.2^S277A^, pYES2-NRT1.2^S292A^, and pYES2-NRT1.2^S277AS292A^, or with the Ser277 phosphomimic form (pYES2-NRT1.2^S277D^), showed similar growth rates to yeast cells transformed with pYES2-NRT1.2; all of these transformed cells exhibited high growth rates. In contrast, yeast cells transformed with Ser292 phosphomimic forms (pYES2-NRT1.2^S292D^, and pYES2-NRT1.2^S277DS292D^) had low growth rates, similar to that of pYES2-NRT1.2+CEPR2 (Fig. 4A, Supplementary Fig. 4C). These results indicated that nonphosphorylated NRT1.2 at Ser292 (NRT1.2^S292A^) might exhibit ABA import activity in yeast cells, and that CEPR2 inhibits the ABA-import activity of NRT1.2 possibly via phosphorylation at Ser292.

**Figure 4.**
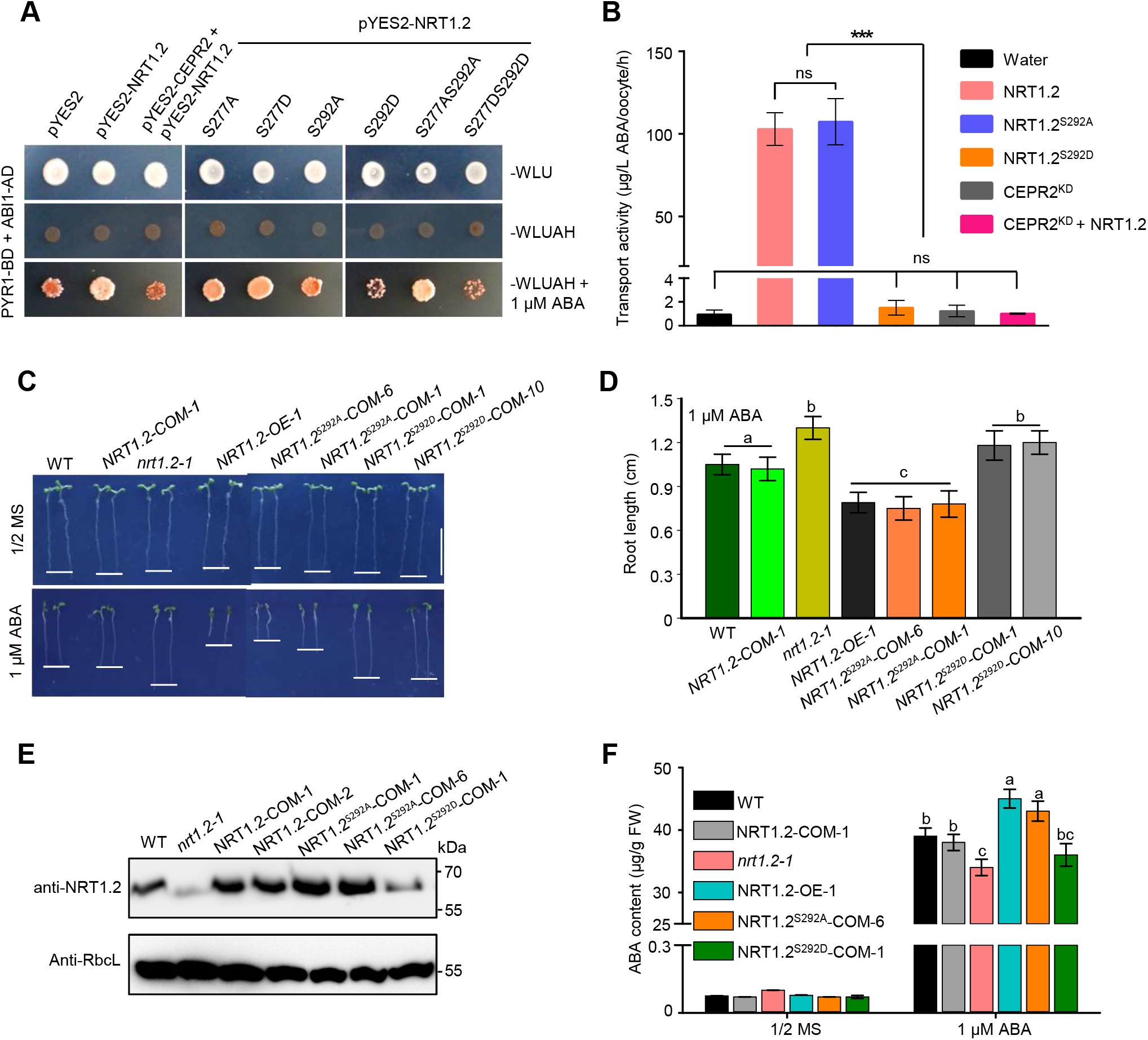
Phosphorylated NRT1.2 was degradated and eliminated the NRT1.2-mediated ABA import. (**A**) Y2H assays showing the ABA transporter activity of NRT1.2. pYES2-NRT1.2 represents normal NRT1.2. pYES2-NRT1.2^S277A^, -NRT1.2^S277D^, -NRT1.2^S292A^, -NRT1.2^S292D^, -NRT1.2^S277AS292A^, and -NRT1.2^S277DS292D^ represent nonphosphorylated and phosphorylated NRT1.2 obtained by point mutant at Ser277 and Ser292 to alanine (A) and aspartate (D), respectively. The pYES2 empty vector was used as a control. (**B**) ABA-import activity in *Xenopus oocytes* expressing NRT1.2, NRT1.2S292A, NRT1.2S292D, CEPR2KD, or NRT1.2 plus CEPR2KD. *X. oocytes* injected with water were used as the negative control. Error bars indicate SEM (N = 3). Statistically significant differences were identified between pairs of measurements using Student’s *t* test (***P < 0.001; ns, not significant). (**C**) The phenotypes of various seedlings grown on 1/2 MS or 1 μM ABA for 7 d: WT, *nrt1.2-1*, NRT1.2 complementary (NRT1.2-COM-1), NRT1.2 overexpression (NRT1.2-OE-1), NRT1.2^S292A^ and NRT1.2^S292D^ complementary (NRT1.2^S292A^-COM, NRT1.2^S292D^-COM). Error bars indicate SEM (N = 3). (**D**) Lengths of the primary roots of the seedlings shown in (C). Error bars indicate SEM (N = 3). Bars labeled with different lowercase letters are significantly different from one another (P < 0.05; one way ANOVA). (**E**) NRT1.2 protein levels in WT, nrt1.2-1, NRT1.2-COM-1, NRT1.2^S292A^-COM, and NRT1.2^S292D^-COM seedlings grown on 1/2 MS for 7 d, as detected using western blots. RbcL was used as the loading control. (**F**) ABA contents in WT, *nrt1.2-1*, NRT1.2-COM-1, NRT1.2^S292A^-COM, and NRT1.2^S292D^-COM seedlings grown on 1/2 MS for 7 d and then treated with ABA for 6 h for HPLC. Error bars indicate SEM (N = 3). Bars labeled with different lowercase letters are significantly different from one another (P < 0.05; one way ANOVA).

We then compared ABA import activity in *Xenopus* oocytes with respect to ABA content. We found that *Xenopus* oocytes expressing NRT1.2 and NRT1.2^S292A^ had similar levels of ABA, which were significantly higher than the negative control *Xenopus* oocytes (injected with water; Fig. 4B). The levels of ABA in *Xenopus* oocytes expressing NRT1.2^S292D^, CEPR2, or CEPR2+NRT1.2 did not differ significantly from the negative control. These results further indicated that NRT1.2^S292A^, but not NRT1.2^S292D^, imports ABA, and that CEPR2 mediates the phosphorylation of NRT1.2 in *Xenopus* oocytes.

Moreover, NRT1.2, NRT1.2^S292A^, and NRT1.2^S292D^ were separately introduced into *nrt1.2-1* mutants to obtain complementary plants (designated NRT1.2-COMs, NRT1.2^S292A^-COMs and NRT1.2^S292D^-COMs, respectively) and verified by RT-PCR (Supplementary Fig. 5A). In the absence of ABA, the growth rates and phenotypes of all transgenic plants, irrespective of genotype, were similar to those of the WT (Fig. 4C). When exposed to ABA, the root growth of the NRT1.2-COM plants was similar to that of the WT. That is, the high root growth of the *nrt1.2-1* plants and the low root growth of NRT1.2-OE-1 plants were both reversed. The root growth of the NRT1.2^S292A^-COM plants was similar to that of the NRT1.2-OE-1 plants, but the root growth of the NRT1.2^S292D^-COM plants were similar to that of the *nrt1.2-1* plants (Fig. 4C and D). NRT1.2 protein levels in NRT1.2^S292A^-COM plants were higher than those in the WT, similar to that in NRT1.2-COM plants, all were higher than WT, while those in NRT1.2 ^S292D^-COM plants showed similar level to those in the *nrt1.2-1* plants (Fig. 4E). Furthermore, ABA levels in the NRT1.2^S292A^-COM plants and the NRT1.2-OE-1 plants were higher than those in the WT, while ABA levels in the NRT1.2^S292D^-COM plants and the *nrt1.2-1* plants were lower than those in the WT (Fig. 5F). Thus, these results indicated that NRT1.2^S292A^ is stable and effectively imports ABA *in planta*.

**Figure 5.**
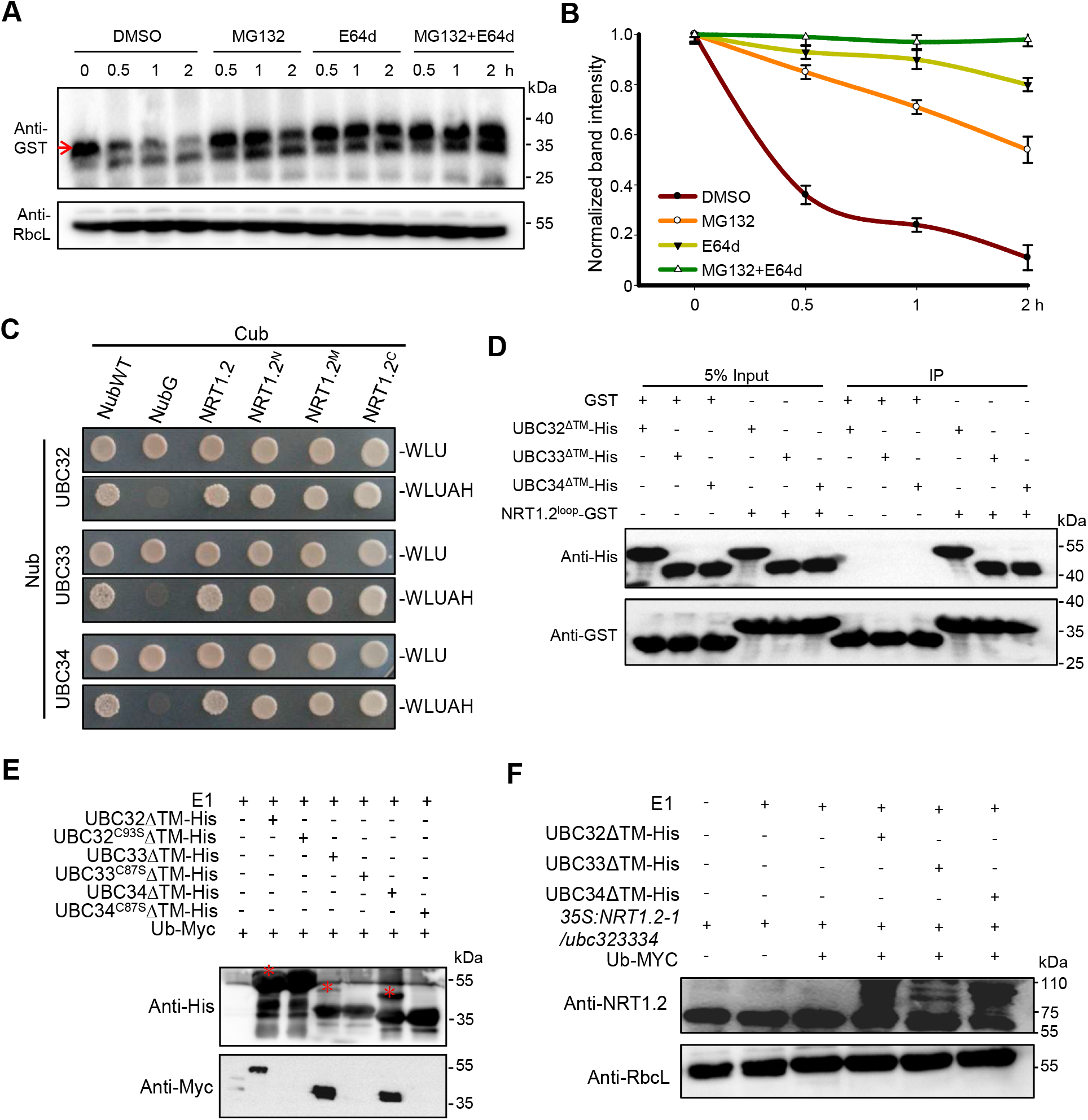
Group XIV ubiquitin-conjugating enzymes UBC32, UBC33 and UBC34 interact with and mediate the degradation of NRT1.2. **(A)** The cell-free degradation of the NRT1.2^loop^-GST protein in the soluble protein extracts from 7-d-old WT seedlings grown on 1/2 MS medium with or without 50 μM MG132 (an inhibitor of the 26S proteasome degradation pathway), and/or 50 μM E64d (an inhibitor of the vacuolar degradation pathway) for 0, 0.5, 1, and 2 h. NRT1.2loop-GST protein levels were quantified using western blots with anti-GST antibody (indicated by red arrows). RbcL was used as the loading control. **(B)** Normalized plot of the NRT1.2 contents, based on the band intensities shown in (A); band intensities were quantified using ImageJ as described in methods. Error bars indicate SEM (N = 3). **(C)** MbSUS assays showing that UBC32, UBC33, and UBC34 interact with the NRT1.2 N-terminal (NRT1.2N, 1–231 aa), loop region (NRT1.2M, 232–346 aa) and the C terminus of NRT1.2 (NRT1.2C, 347-585 aa). NubWT and NubG are treated as positive and negative controls, respectively. **(D)** Pulldown assays using purified GST, NRT1.2loop-GST, UBC32ΔTM-His, UBC33ΔTM-His and UBC34ΔTM-His expressed in *E.coli*. ΔTM indicates the deletion of the transmembrane regions of UBC32, UBC33, and UBC34. **(E)** In vitro assays of UBC32ΔTM-His, UBC33ΔTM-His, and UBC34ΔTM-His purified from *E.coli* to test for self-ubiquitination. Note that contructs with point mutants (UBC32C93SΔTM-His, UBC33C87SΔTM-His and UBC34C87SΔTM-His) do not self-ubiquitinate. **(F)** Semi *in vivo* ubiquitination assay, using 35S:NRT1.2-1/ubc323334 and UBC32ΔTM-His, UBC33ΔTM-His, UBC34ΔTM-His (purified from *E*.coli) to test the ubiquitination of NRT1.2 by UBC32, UBC33, and UBC34.

### UBC32, UBC33, and UBC34 mediate the degradation of NRT1.2

To further investigate the NRT1.2 degradation pathway, protein extracts from 7-d-old CEPR2-OE-9 seedlings were treated with 50 μM MG132 (an inhibitor of the 26S proteasome degradation pathway), 50 μM E64d (an inhibitor of the vacuolar degradation pathway), or both prior to incubation with purified NRT1.2^loop^-GST. After incubation, NRT1.2 degradation was significantly inhibited by treatment with MG132 and/or E64d, but not by treatment with dimethyl sulfoxide (DMSO, the control; Fig. 5A and B). For example, after 2 h incubation of NRT1.2^loop^-GST with protein extracts from 7-d-old CEPR2-OE-9 seedlings treated with DMSO, 11% of NRT1.2^loop^-GST remained; in contrast, after 2 h incubation of NRT1.2^loop^-GST with protein extracts from 7-d-old CEPR2-OE-9 seedlings treated with MG132, E64d or both, 58%, 80% and 99% NRT1.2^loop^-GST remained (Fig. 5B). These results suggested that NRT1.2 degradation is mediated by both the 26S proteasome and the vacuolar degradation pathways.

To identify the components that mediate the degradation of NRT1.2, we searched the BioGRID web site (https://thebiogrid.org/), and found UBIQUITIN-CONJUGATING ENZYME 34 (UBC34) among the interacting proteins (Xu et al., 2020). UBC34, which is an E2 ligase with ubiquitination activity, is highly similar to UBC32 and UBC33 (Cui et al., 2012a; Cui et al., 2012b). Our results showed that NRT1.2 interacted with UBC32, UBC33 and UBC34 in yeast (Fig. 5C). Pulldown assays using the NRT1.2 and transmembrane-region-deleted UBCs (Supplementary Fig. 6A) further confirmed the interaction between NRT1.2^loop^ and the three UBCs (Fig. 5D).

The UBC-mediated ubquitination of NRT1.2 was further investigated. *In vitro* ubiquitination analysis indicated that UBC32, UBC33, and UBC34 self-ubiquitinated (Fig. 5E), confirming that these UBCs possessed ubiquitination abilities. However, when point-mutated forms of UBC32 (C93S), UBC33 (C87S), or UBC34 (C87S) were used, neither ubiquitin nor ubiquitination were detected (Fig. 5E), suggesting that Cys93 (in UBC32) and Cys87 (in UBC33 and UBC34) played important roles in ubiquitination. Next, the UBC-dependent ubiquitination of NRT1.2 was tested using a semi-*in vitro* assay. In this assay, protein extracts from a triple mutant of UBC32, UBC33, and UBC34 (*ubc323334*) overexpressing NRT1.2 were incubated with purified UBC32ΔTM-His, UBC33ΔTM-His, or UBC34ΔTM-His. Ubiquitinated NRT1.2 was only detected in the presence of UBC32, UBC33, or UBC34 (Fig. 5F), suggesting that UBC32, UBC33, and UBC34 mediated the ubiquitination of NRT1.2.

Finally, we generated *NRT1.2-OE-1/ubc323334* and *nrt1.2ubc323334* mutant plants. We overexpressed *NRT1.2* in *ubc323334* mutant plants and crossed *nrt1.2* with *ubc323334* mutant plants. The obtained plants were confirmed using RT-PCR (Supplementary Fig. 5B and C). Genetic analysis showed that, during seed germination, the *NRT1.2-OE-1/ubc323334* plants had similar phenotype to the *ubc323334* and *NRT1.2-OE-1* plants. These transgenic plants were all more sensitive than the WT plants to ABA. In contrast, the *nrt1.2ubc323334* mutant had a similar phenotype to that of *nrt1.2-1* mutant, and was less sensitive to ABA than the WT (Fig. 6A and B). When exposed to ABA, the root lengths of the *NRT1.2-OE-1/ubc323334, ubc323334* and *NRT1.2-OE-1* plants were similar to each other and shorter than that of the WT; the root lengths of the *nrt1.2ubc323334* and *nrt1.2-1* mutants were also similar and longer than that of the WT (Fig. 6C and D). Under normal conditions, the NRT1.2 protein content was obviously greater in the *ubc323334* mutant than in the WT plants. However, NRT1.2 protein content in the *ubc323334* mutant was similar to that in the WT plants in the presence of ABA (Fig. 6E and F). These data indicated that UBC32, UBC33, and UBC34 act epistatically and negatively on NRT1.2 via ubiquitination modification.

**Figure 6.**
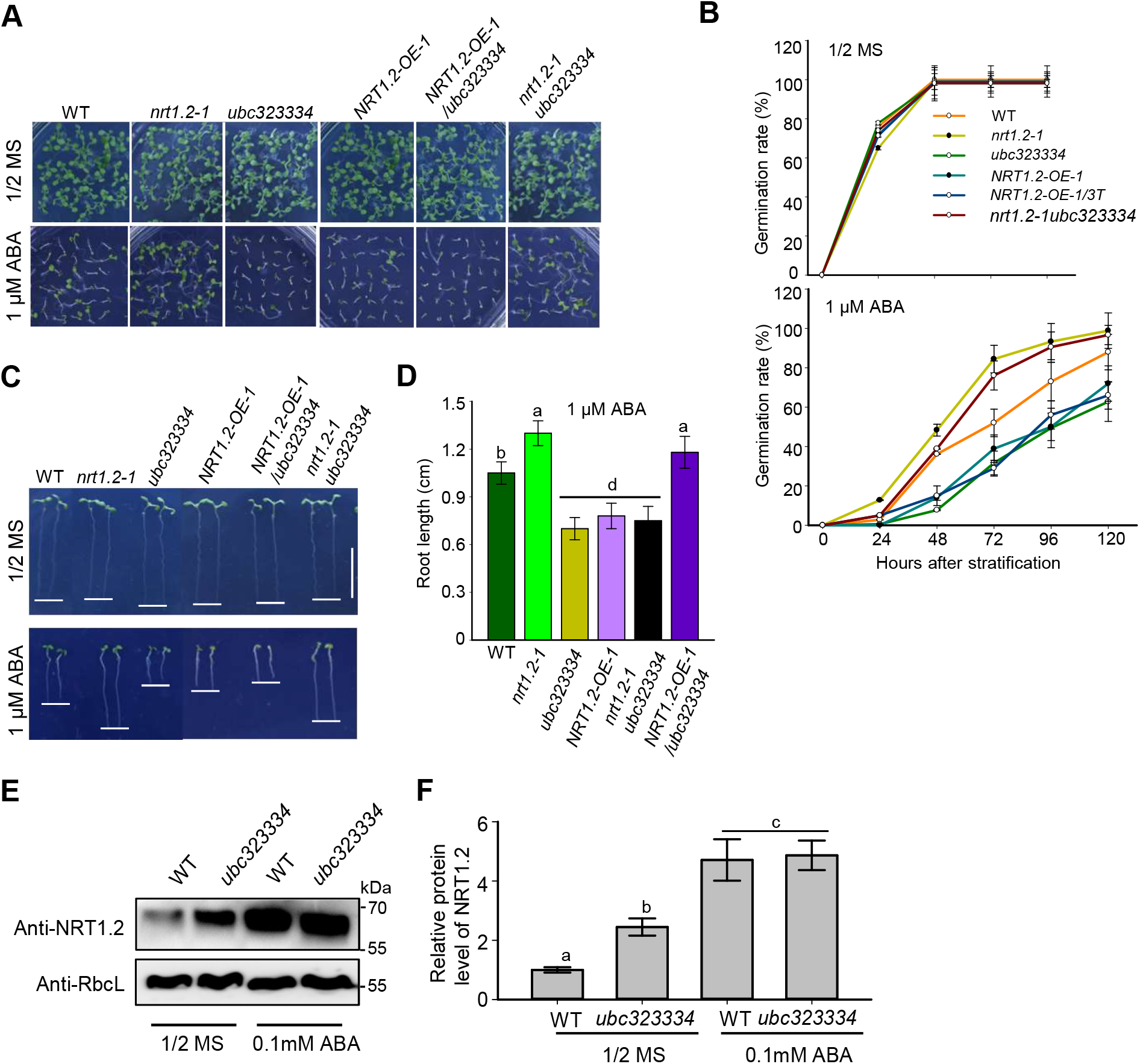
UBC32, UBC33, and UBC34 act epistatically on NRT1.2. **(A)** The germination phenotype of WT, *nrt1.2-1, ubc323334*, NRT1.2-OE-1, NRT1.2-OE-1/*ubc323334*, and *nrt1.2-1 ubc323334* plants grown on 1/2 MS with or without 1 μM ABA for 10 d. **(B)** The germination rates of the plants shown in (A).The upper panel shown rates on 1/2 MS of rate, and the lower panel shown rates on 1 μM ABA. Error bars indicate SEM (N = 3). **(C)** Phenotypes of seedlings with different genotypes grown on 1/2 MS with or without 1 μM ABA for 7 d. **(D)** Lengths of the primary roots of the seedlings shown in (C). Error bars indicate SEM (N = 3). Bars labeled with different lowercase letters are significantly different from one another (P < 0.05; one way ANOVA). **(E)** NRT1.2 protein levels in WT and *ubc323334* seedlings grown on 1/2 MS for 7 d before treatment with or without 0.1mM ABA for 6 h. Protein levels were measured using western blots. RbcL was used as the loading control. **(F)** Normalized plot of the NRT1.2 degradation shown in (E), based on the band intensities: band intensities were quantified using ImageJ as described in methods. Error bars indicate SEM (N = 3).

## Discussion

ABA transporters play critical roles in the systematic ABA response in plants (Kuromori et al., 2018). Although, several ABA transporters have been identified (Huang et al., 1999; Kang et al., 2015; Kanno et al., 2012; Kuromori et al., 2010; Zhang et al., 2014), the mechanisms regulating ABA transporters remain largely unknown. The identification of CEPR2 as well as UBC32, UBC33, and UBC34 as regulators of NRT1.2 in this study helps to clarify the molecular mechanisms modulating ABA import.

In most cases, protein phosphorylation rapidly and efficiently modifies target proteins in response to environmental stimuli (Grondin et al., 2015). Studies have shown that transporter activity can be regulated via the phosphorylation of proteins, such as SOS1 (Quintero et al., 2011) and PIP2.1 (Grondin et al., 2015). Our results demonstrated that the phosphorylation of NRT1.2 is partially mediated by CEPR2. The phosphorylation of NRT1.2 promotes its degradation, leading to the suppression of ABA import and the ABA response (Fig. 2–4). The differential phosphoproteomic analysis of CEPR2-OE-9 versus the WT showed that NRT1.2 phosphorylation was affected by CEPR2 (Supplementary Excel 1). One of the two identified phosphorylation sites was crucial for NRT1.2 stability due to its inhibition of degradation. This site was also identified as significant for CEPR2-mediated phosphorylation based on the lack of phosphorylation at NRT1.2^S292A^ (Fig. 3). ABA strongly inhibited CEPR2-mediated NRT1.2 degradation (Fig. 3). Recent reports have indicated that CEPR2 has similar effects on ABA receptors, such as PYL4 (Yu et al., 2019). However, non-phosphorylated NRT1.2 was stable and was able to import ABA into cells (Fig. 4). The increased concentrations of intracellular ABA were recognized by the non-phosphorylated PYLs, eventually stimulating ABA signaling. Therefore, CEPR2-mediated phosphorylation represents an important mechanism that maintains the transition between the ABA response and plant growth (Fig. 7).

**Figure 7.**
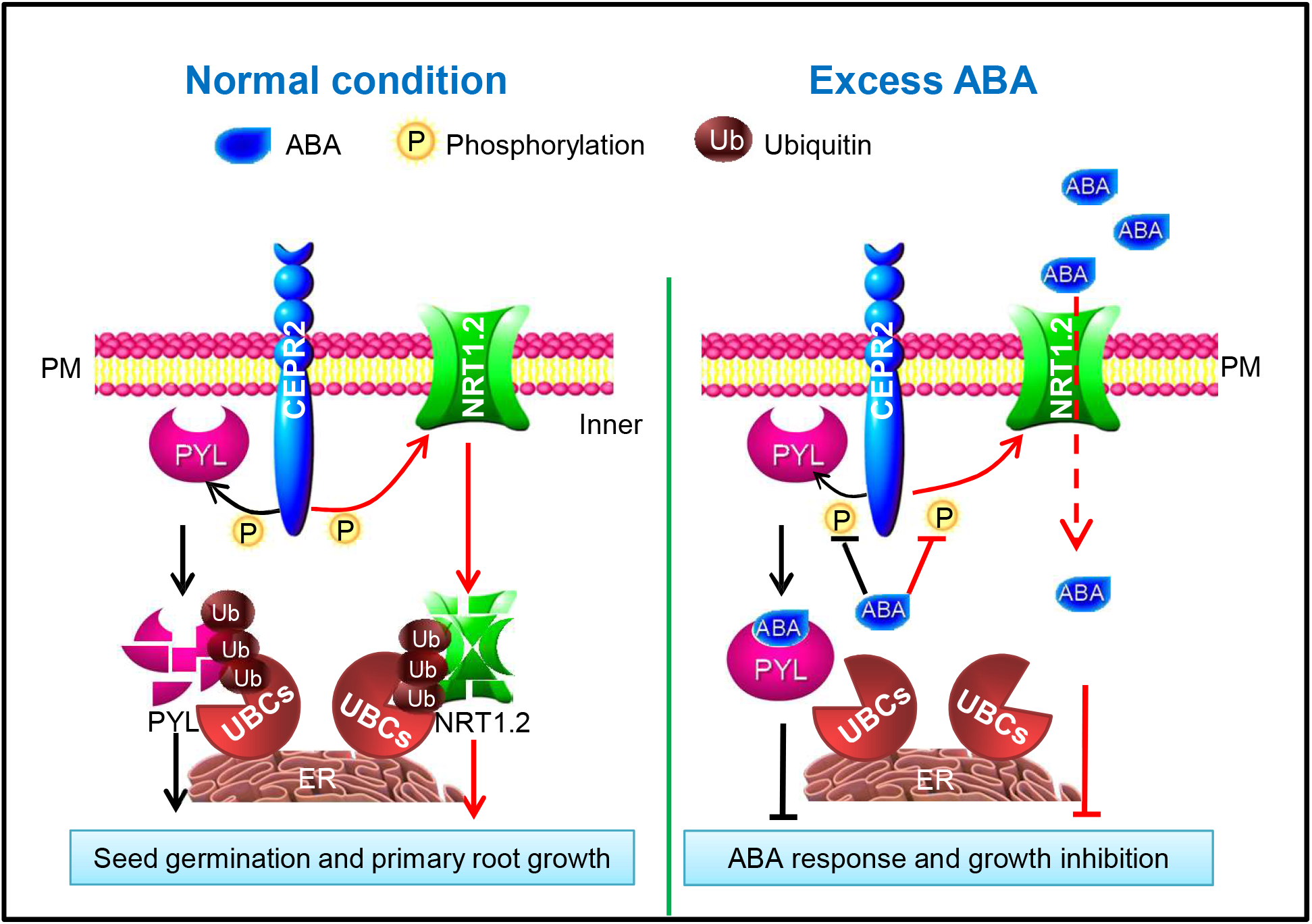
Phosphorylation and ubiquitination of NRT1.2 by CEPR2 regulate ABA influx and plant growth in *Arabidopsis*. Under normal conditions, CEPR2 first phosphorylates NRT1.2 and PYLs at the plasma membrane. The phosphorylated NRT1.2 and PYLs might then be translocated to ER, where they are ubiquitinated by UBC32, UBC33, and UBC34. Finally, the ubiquitinated NRT1.2 and PYLs are degraded by 26S proteasome and the vacuolar degradation pathway. Thus, the seed germination and primary roots grow normally. However, ABA inhibits the phosphorylation of NRT1.2 and PYLs by CEPR2, and stabilizes these proteins, stimulating NRT1.2 to import ABA into the cells, where ABA is bound by PYLs to cause the ABA response, which leads to the inhibition of seed germination and primary root growth.

Further investigation indicated that the degradation of NRT1.2 was significantly inhibited by MG132 alone, E64d alone, and MG132 plus E64D (Fig. 5A and B). This suggested that NRT1.2 degradation was mediated both by the 26S proteasome and by the vacuoles. Y2H and pulldown assays showed that AtUBC32, AtUBC33, and AtUBC34 interacted with NRT1.2 (Fig. 5C and D), ubiquitinated as well as themselves (Fig. 5E and F), and epistatically and negatively regulated NRT1.2 in response to ABA (Fig. 6). These results demonstrated that these three UBCs mediate the ubiquitination and degradation of NRT1.2. UBC32 is a functional E2 that negatively regulates salt tolerance in *Arabidopsis* (Cui et al., 2012b). Subcellular localization analysis indicated that AtUBC32 is located in the membrane of endoplasmic reticulum (ER), and is an active ER-associated degradation (ERAD) component, playing significant roles in the plant ERAD system (Cui et al., 2012a). The ERAD process consists of substrate recognition, targeting, retrotranslocation, polyubiquitination, and degradation by the 26S proteasome (Cui et al., 2012a). UBC32, UBC33, and UBC34 are homologous in Arabidopsis (Cui et al., 2012b). UBC34 from *Populus tomentosa* (PtoUBC34) was also localized in the ER membrane, where it interacts with the transcriptional repressors PtoMYB221 and PtoMYB156 (Zheng et al., 2019). This specific interaction allowed the translocation of PtoMYB221 and PtoMYB156 to the ER, and reduced their suppression of genes involved in lignin biosynthesis (Zheng et al., 2019). The translocation of PtoMYB221 and PtoMYB156 to the ER by the ER-localized UBC34 suggests that PtoMYB221 and PtoMYB156 may be degraded via the ERAD pathway. Furthermore, AtUBC34 is found to trigger the turnover of SUCROSE TRANSPORTER 2 (SUC2) in response to light and ubiquitinates SUC2 *in vitro* (Xu et al., 2020). Recently, VPS23a, an E2-like protein, was identified as a key component of the endosomal sorting complex required for transports (ESCRTs) machinery I and promoted vacuolar degradation by PYL4 (Xia et al., 2020; Yu et al., 2016). These data suggested that NRT1.2 might be ubiquitinated by AtUBC32, AtUBC33, and AtUBC34 at ER membrane, and then degraded by the 26S proteasome and vacuolar degradation pathways.

However, almost half of NRT1.2 was nonetheless degraded in the *cepr2-1* mutant (Fig. 3A and B), and CEPR2 did not phosphorylate Ser277 in NRT1.2 (Fig. 3C and D). In addition, the phosphorylation of NRT1.2 at Ser277 might not affect its ABA import activity, as indicated by the ABA transporting assay in yeast cells (Fig. 4A). Finally, based on the levels of the NRT1.2 protein, NRT1.2 degradation also occurred in the WT and *ubc323334* plants under normal conditions, as compared to the ABA treatment (Fig. 6E and F). These data suggested that other components mediate the phosphorylation and degradation of NRT1.2 under normal conditions, and that NRT1.2 might have functions beyond ABA import. To fully characterize the regulation of NRT1.2 phosphorylation, degradation, and function, the proteins that interact with NRT1.2 must be more fully identified and characterized.

In summary, our results indicated that the phosphorylation and ubiquitination of NRT1.2 by CEPR2 and UBCs regulate ABA influx and plant growth in *Arabidopsis* (Fig. 7). That is, under normal condition, CEPR2 first phosphorylates NRT1.2 and PYLs at the plasma membrane. The phosphorylated NRT1.2 and PYLs might then be translocated to the ER, where they are ubiquitinated by UBC32, UBC33, and UBC34. Finally, the ubiquitinated NRT1.2 and PYLs are degraded by 26S proteasome and the vacuolar degradation pathway, allowing seed germination and normal primary root growth. However, ABA inhibits the phosphorylation of NRT1.2 and PYLs by CEPR2, and stabilizes these proteins, stimulating NRT1.2 to import ABA into the cells, where ABA is bound by PYLs to cause the ABA response, leading to the inhibition of seed germination and primary root growth.

## Materials and Methods

### Plant materials and growth conditions

*Arabidopsis thaliana* (L.) Heynh. cv. ‘Columbia-0’ was used as the WT. The T-DNA insertion lines, SALK_072696 (*nrt1.2-1*), SALK_011182 (*nrt1.2-2*) in the Col-0 background were obtained from the *Arabidopsis* Biological Resource Center (http://www.arabidopsis.org). The *cepr2-1* T-DNA insertion mutant and the CEPR2-overexpressing line CEPR2-OE-9 have previously been characterized (Yu et al., 2019). The homozygous single and double mutants, *cepr2-1, nrt1.2-1*, and *cepr2-1nrt1.2-1* were verified using RT-PCR with the primers listed in Supplementary Table 2. The *nrt1.2-1* complementary seedlings with a point mutation at Ser^292^ to alanine and to aspartate were designated NRT1.2^S292A^-COM and NRT1.2^S292D^-COM seedlings. NRT1.2 overexpression seedlings, NRT1.2-COM seedlings, NRT1.2^S292A^-COM seedlings, and NRT1.2^S292D^-COM seedlings were selected on 1/2 Murashige and Skoog (MS) medium (1.5% sucrose and 0.85% agar) supplemented with 50 mg/L kanamycin. *NRT1.2* expression levels in the *NRT1.2* transgenic seedlings were confirmed with RT-PCR using the primers listed in Supplementary Table 2. All selected *Arabidopsis* plants were grown in a greenhouse under a 16 h light/8 h dark cycle at 23°C. The light was 9600 LUX.

### Phosphoproteome analysis

Protein extraction and phosphoproteome analysis were performed as previously described (Pi et al., 2016). We extracted total proteins from CEPR2-OE-9 and WT seedlings grown in 1/2 MS for 7 d using the TCA/Acetone extraction method. Phosphoproteome analysis was performed using capitalBio technology (Beijing, China). The mass spectrometry data were deposited in the ProteomeXchange Consortium via the PRIDE partner repository (dataset identifier PXD021458; http://www.ebi.ac.uk/pride).

### MbSUS, BiFC, and LCI assays

The MbSUS assay was performed as described previously (Obrdlik et al., 2004); detailed descriptions of associated experimental principles and methods were given in Yu et al. (2019). In brief, for Nub fusions, PCR products were cloned and transformed with pNXgate, cleaved with *Eco*RI and *Sma*I, and transformed into the THY.AP5 yeast strain. Strains were selected on synthetic dropout (SD) medium lacking tryptophan (W) and uracil (U). For Cub fusions, PCR products were coloned and transformed with pMetYCgate, cleaved with *Pst*I/*Hind*III, and transformed (together with PCR products) into the THY.AP4 yeast strain. Transformants were further selected on SD medium lacking leucine (L). Clones from each transformant were incubated on SD media lacking leucine, tryptophan, and uracil (SD-Trp/-Leu/-Ura; SD-WLU) at 30°C for three days. To detect protein interactions, colonies were spotted onto control media (SD-WLU), as well as onto selection media lacking leucine, tryptophan, uracil, adenine, and histidine (SD-Trp/-Leu/-Ura/-Ade/-His; SD-WLUAH). Colony growth was monitored for 3-6 days.

Luciferase complementary imaging (LCI) assays were performed as previously described (Chen et al., 2008; Xu et al., 2018). In brief, the full-length CDS of *CEPR2* was cloned into the pCAMBIA1300-nLUC (nLUC) vector to generate the CEPR2-nLUC construct, and the full-length CDS of *NRT1.2* was cloned into the pCAMBIA1300-cLUC (cLUC) vector to generate the NRT1.2-cLUC construct. Next, every 1 mL of *Agrobacterium tumefaciens* cells harboring CEPR2-nLUC, NRT1.2-cLUC, and nLUC were mixed to obtain the following combinations, each with a final optical density at 600 nm (OD_600_) of 1.0: CEPR2-nLUC + cLUC, NRT1.2-cLUC + nLUC, and CEPR2-nLUC + NRT1.2-cLUC. Each of these combinations of *A. tumefaciens* cells were separately infiltrated into *N. benthamiana* leaves. *N. benthamiana* plants were grown at 26°C for 60 h. Five minutes before detection, 0.2 mM luciferin (Promega, Madison, WI, USA) was sprayed on the treated leaves, and luciferase activity was then measured using a cooled charge-coupled device (Lumina II, Waltham, MA, USA).

BiFC assays were performed as described previously (Waadt and Kudla, 2008). In brief, the full-length CDS of *CEPR2* (without the stop codon) was cloned into the pSPYNE-35S vector to generate the CEPR2-YFP^N^ construct, and the full-length CDS of *NRT1.2* (without the stop codon) was cloned into the pSPYCE-35S vector to obtain the NRT1.2-YFP^C^ construct. *A.tumefaciens* cells harboring CEPR2-YFP^N^ and NRT1.2-YFP^C^ were mixed with 10 mL MMA medium (10 mM MgCl_2_, 50 mM MES, and 20 μM acetosyringone, pH 5.6) to yield a final optical density at 600 nm (OD_600_) of 1.0. The cell mixtures were then injected into *N.benthamiana* leaves by gently pressing a disposable syringe to the abaxial surface of a fully-expanded leaf (approximate width of 3 cm at the midpoint). Plants were grown at 26°C for 36–60 h, and then YFP signals in the leaves were detected at 488 nm using a Confocal Laser Scanning Microscope LSM51 (Zeiss, Germany). Here, the amphiphilic styryl dye FM4-64 (N-(3-triethylammomiumpropyl) 4-(p-diethylaminophenylhexa-trienyl)) was used as a plasma membrane marker.

### Pulldown assay

To characterize the interactions between CEPR2, UBCs, and NRT1.2, the CDS encoding the cytosol loop region (232–346 aa) of NRT1.2 was fused with GST in the pGEX-4T-1 vector to obtain the pGEX-4T-1-GST-NRT1.2 (NRT1.2-GST) construct; the full length CDSs of the UBCs (without transmembrane domains) were fused with the His tag in the pET30a-His vector to obtain the pET30a-His-UBC32ΔTM-His, pET30a-His-UBC33ΔTM-His, and pET30a-His-UBC34ΔTM constructs; the kinase domain (642-977 aa) of CEPR2 was fused with His tag in pET30a-His vector to obtain pET30a-His-CEPR2^KD^-His (CEPR2^KD^-His) construct. The obtained NRT1.2-GST and CEPR2^KD^-His constructs were transformed into competent cells of the *E. coli* Rosetta strain. The transformed cells were cultured in 500 mL Luria-Bertani medium at 37°C to an OD_600_ of 1.0. Protein expression was then induced with 0.8 mM Isopropyl β-D-Thiogalactoside (IPTG) for 12 h at 16°C. Next, the *E. coli* cells were obtained by centrifugation at 6000 rpm for 5 min at 4°C. The pellet was resuspended in 5 mL ddH_2_O. Lysates were obtained using ultrasonication (JY92-II, Scientz Biotechnology Co., Ltd, Ningbo, China) with the following parameters: operating power, 300 w; working time, 10 s; interval time, 5 s; and cycles, 30. Lysates were clarified by centrifugation at 8000 rpm for 10 min at 4°C. Then, the CEPR2^KD^-His protein was purified using the His-Tagged Protein Purification Kit (CWBIO, Beijing, China), and NRT1.2-GST was purified using Pierce Glutathione Spin Columns (Thermo, Waltham, MA, USA). In the pulldown assay, 50 μg NRT1.2^loop^-GST and 50 μg CEPR2^KD^-His were incubated in 1 mL binding buffer (50 mM Tris-HCl, 150 mM NaCl, pH 8.0) at 4°C for 2 h with constant slight shaking. After incubation, GST proteins were purified with Pierce Glutathione Spin Columns, eluted, and analyzed using anti-His antibodies (CWBIO, Beijing, China). The primers used in this experiment are listed in Supplementary Table 2.

### RNA extraction, RT-PCR, and quantitative RT-PCR (qRT-PCR)

Total RNAs from 7-d-old seedlings grown on 1/2 MS with or without 1 μM ABA were extracted using TRIzol (Invitrogen, Carlsbad, CA, USA) or Universal Plant Total RNA Extraction Kits (Spin-column)-I (BioTeke, Beijing, China). The cDNA used for qRT-PCR were synthesized using PrimeScript reverse transcriptase with oligo dT primer and Prime Script RT Enzyme MIX I (Takara, Osaka, Japan). qRT-PCRs was performed using the ChamQ SYBR Color qPCR Master Mix (Q411, Vazyme, Nanjing, China) on a Bio-Rad CFX96 (Bio-Rad, Hercules, CA, USA). *UBC21* and *UBQ10* were used as internal controls for qRT-PCR. The cDNAs used for RT-PCR were synthesized using PrimeScript First-Strand cDNA Synthesis Kits (Takara, Osaka, Japan). RT-PCR cycling conditions were as follows: denaturation at 94°C for 5 min, followed by 23-34 cycles of amplification, and a final elongation at 72°C for 5 min. *EF-1*α was used as the internal control for RT-PCR. The primers used are listed in Supplementary Table 2.

### Germination and root length measurements

Plants with different genotypes were grown under same conditions in the greenhouse, and all seeds were harvested simultaneously. Germination assays were performed as previously described (Chen et al., 2014). In brief, the seeds were surface-sterilized and incubated on 1/2 MS medium with or without 1 μM ABA in the dark at 4°C for three days. Germination was assessed every 12 h; germination was defined as the emergence of the radicle through the seed coat. At the end of the three days, germination rates were calculated. To assess growth, sterilized seeds were grown vertically on 1/2 MS medium with or without 1 μM ABA for seven days. The root lengths of at least 20 seedlings per line (WT, mutant, and transgenic) were measured using a ruler, and mean root length was calculated for each line.

### Cell-free degradation

To investigate the effects of CEPR2 on NRT1.2, NRT1.2-GST protein levels were detected after incubation with total protein extracts from CEPR2-OE-9, *cepr2-1*, or WT seedlings grown on 1/2 MS medium with or without 1 μM ABA. Total proteins were extracted from 0.4 g samples of 7-d-old seedlings of all genotypes using 600 μL extraction buffer (5 mM DTT, 10 mM NaCl, 25 mM Tris-HCl [pH 7.5], 10 mM ATP, 4 mM PMSF, and 10 mM MgCl_2_). The crude extracts were held on ice (4°C) for 30 min and centrifuged twice at 12,000 rpm for 10 min at 4°C. After centrifugation, the supernatants were collected. About 0.5 mg of the purified NRT1.2-GST was incubated with the total protein extracts at 22°C for 0, 10, 30, or 60 min. At each time point, 20 μL of each solution was transferred to a new centrifuge tube, combined with 5 μL of 5× loading buffer, and boiled for 5 min to terminate the reaction. The abundance of the NRT1.2 protein in each reaction was detected using anti-GST antibodies. Spot densitometry was measured using ImageJ v1.36 (http://rsb.info.nih.gov/ij/). To investigate the NRT1.2 degradation pathway, protein extracts with 100 μM MG132 (an inhibitor of the 26S proteasome pathway) and/or 100 μM E64d (an inhibitor of the vacuolar degradation pathway) were incubated with purified NRT1.2^loop^-GST fusion proteins. Protein levels were detected using anti-GST antibodies. ImageJ v1.36 was used to quantify the intensity of each protein band.

### *In vitro* kinase assay

To investigate possible sites at which CEPR2 phophorylated NRT1.2, *in vitro* kinase assays were carried out. The CDSs encoding the mutant loop region (232–346 aa) of NRT1.2 in which Ser^277^ or Ser^292^ were point mutated to Alanine (A) or aspartate (D), were each fused with the GST tag in the pGEX-4T-1 vector to obtain the pGEX-4T-1-GST-NRT1.2^loopS277A/D^ (NRT1.2^loopS277A/D^-GST) and pGEX-4T-1-GST-NRT1.2^loopS292A/D^ (NRT1.2^loopS292A/D^-GST) constructs. The transformation, induction and purification of these fusion proteins were performed as described above. CEPR2-GFP was purified using affinity chromatography. *In vitro* kinase assays were carried out using purified CEPR2-GFP by and NRT1.2^loop^-GST, NRT1.2^loopS277A^-GST, or NRT1.2^loopS292A^-GST. We combined 50 μg of NRT1.2^loop^-GST, NRT1.2^loopS277A^-GST, or NRT1.2^loopS292A^-GST with 0.5 μg CEPR2-GFP in 50 μL of reaction buffer (25 mM HEPES, pH 7.2, 1 mM DTT, 50 mM NaCl, 2 mM EGTA, 5 mM MgSO_4_, and 50 μM ATP). The reaction mixtures were incubated at 30°C for 0-120 min and the reaction was terminated by adding 5× loading buffer. The unincubated reaction mixture (at 0 min) was used as the negative control. Proteins were then fractionated on sodium dodecyl sulfate polyacrylamide gel electrophoresis (SDS-PAGE) and Mn^2+^-phos-tag-PAGE (50 μM phos-tag and 100 μM Mn^2+^). After 30 min incubation at 37°C, 0.01 U/μL Calf Intestinal Alkaline Phosphatase (CIAP, Promega, Madison, WI, USA) was added to the reaction buffer, and the mixture was incubated for another 30 min at 37°C to remove the phosphoryl group(s) of NRT1.2^loop^. To determing the effects of ABA on the phosphorylation of NRT1.2, *in vitro* kinase assays were performed using purified CEPR2^KD^-His and NRT1.2-GST with or without 10 μM ABA in the presence of 100 μM MG132 and 100 μM E64d. Primers used in this experiment are listed in Supplementary Table 2.

### Analysis of ABA-import activity in a yeast system

The CDSs of PYR1 and ABI1 were amplified using PCR, and inserted into the pGBKT7 and pGADT7 plasmids, respectively. Then, the plasmids were co-transformed into *Saccharomyces cerevisiae* Yeast Gold cells. Successfully transformed colonies were identified on SD media lacking Leu and Trp (SD-WL). To detect ABA import activity levels, pYES2 plasmids or plasmids containing the full-length CDS of NRT1.2, NRT1.2^S292A^, or NRT1.2^S292D^ were separately transformed into yeast cells. Successfully transformed colonies were identified on SD-WLU media, then transferred to SD-WLUAH in the absence or presence of 1 μM ABA. ABA-import activity was quantified by observating the growth of the transformed cells, because the growth of the transformed yeast cells was depend on ABA levels. Growth was photographed after 2-3 days of culture. Yeast transformants were transferred to 5 mL of SD-WLUAH liquid medium and incubated at 30°C for 8 h in a shaker. The absorbance at 600 nm was measured using a spectrophotometer (T6, Pgeneral, Beijing, China). Twelve replicates were performed for each cotransformantoin, and mean values were calculated. Primers used in this experiment are listed in Supplementary Table 2.

### Quantification of NRT1.2 transporter activity in *Xenopus* oocytes

To further test the effects of CEPR2 phosphorylation on NRT1.2-mediated ABA import, we measured the levels of ABA transport activity associated with NRT1.2, nonphosphomimic-NRT1.2 (NRT1.2^S292A^), and phosphomimic-NRT1.2 (NRT1.2^S292D^) in the *X*.oocytes. The QuikChange Lightning site-directed mutagenesis kit (Agilent Technologies, CA, USA) was used to generate NRT1.2^S292A^ and NRT1.2^S292D^, which were then cloned into the expression vector pT7TS. The pT7TS plasmids were linearized by *Bam*HI, and the circular RNA (cRNA) was transcribed *in-vitro* using an RNA synthesis kit (mMESSAGE mMACHINET7 kit; Ambion). Primers used in these experiments are listed in Supplementary Table 2. *Xenopus* oocytes were isolated in 25 mL of ND96 solution, containing 43 mg collagenase and 12.5 mg trypsin inhibitor, for 1.5 h. After isolation, cells were recovered in 25 mL of ND96 for another 24 h. The *Xenopus oocytes* were then injected with 20 ng of NRT1.2, NRT1.2^S292A^, NRT1.2^S292D^, and/or CEPR2^KD^ cRNA. Injected *Xenopus* oocytes were incubated in MBS with or without 10 μM ABA for 6 h, and then washed five times with MBS. ABA contents in the oocytes were measured using liquid chromatography-tandem mass spectrometry (LC-MS/MS).

### Analysis of NRT1.2 transporter activity *in planta*

To investigate the effects of NRT1.2 phosphorylation on ABA import *in planta*, we first generated transgenic *nrt1.2-1* mutants carrying native promoter-driven normal NRT1.2 (NRT1.2-COMs), NRT1.2 not phosphorylated at Ser^292^ (NRT1.2^S292A^-COMs), or NRT1.2 phosphorylated at Ser^292^ (NRT1.2^S292D^-COMs) as described above. To measure ABA content, various seedlings (i.e. the NRT1.2^S292A^-COM-8, NRT1.2^S292D^-COM-2, and *nrt1.2-1* mutants, NRT1.2-OE-1, and WT) were grown on 1/2 MS with or without 1 μM ABA for seven days. Next, ABA contents in the seedlings were measured as previously described (Xu et al., 2016).

### Ubiquitination assays

The constructs pET30a-His-UBC32ΔTM-His, pET30a-His-UBC33ΔTM-His, and pET30a-His-UBC34ΔTM, as well as the corresponding mutant fusion proteins, were expressed and purified as described above. The thioester assay reaction was performed in a 60 μL reaction volume, containing 50 mM Tris-HCl (pH 7.4), 10 mM MgCl2, 10 mM ATP, 100 ng UBE1 (Boston Biochemicals, Cambridge, MA, USA), 10 μg recombinant, and 10 μg pET30a-His-UBC33ΔTM-His, pET30a-His-UBC32ΔTM-His pET30a-His-UBC34ΔTM, or the corresponding mutant recombinant Ub-Myc (Boston Biochemicals, Cambridge, MA, USA). Reactions were split after incubation for 6-24 hours at 30 °C and terminated using SDS sample buffer without dithiothreitol (DTT). Samples were separated on a 10% SDS-PAGE gel after boiling at 100 °C for 5 min, and then visualized using western blot with anti-His and anti-Myc.

Total membrane proteins isolated from the *35S:NRT1.2-1/ubc323334* mutant were incubated with purified UBCs. Then, immunoprecipitation and immunoblot analysss were performed as described above. The monoclonal NRT1.2 antibody (Sangon Biotech China) was used for detecting ubiquitinated proteins.

## Author contribution

C.W. and C.Z. conceived the original screening and research plans; L.Z., Z.Y., X.Y., Y.M. and Y.R. performed experiments, analyzed the data, made the figures, and wrote the original article; G.Y., S.Z., J.H., and K.Y. provided suggestions; C.W. and C.Z. supervised and complemented the writing. All authors read and approved the final manuscript.

## Acknowledgments

This work was supported by the Major Program of Shandong Province Natural Science Foundation (ZR2018ZB0212), the National Key R&D Program of China (2018YFD1000704, 2018YFD1000700), and the Natural Science Foundation of China (Grant number 31970292, 31570271). We thank Prof. Yi Wang (Chinese Agricultural University) for the measurement of NRT1.2 transporter activity in *Xenopus* oocyte cells, and Prof. Kim Woo Taek for donating the *ubc323334* triple mutant from Yonsei University. We also thank LetPub (https://www.letpub.com/) for its linguistic assistance during the preparation of this manuscript.

## Data availability

All relevant data, vectors, and plant materials that support the findings of this study are available from the corresponding author upon request.

## Supplementary data

**Supplementary Figure 1.** Phosphoproteome analysis and mating-based split ubiquitin system assay (MbSUS) assay to detect the interaction between CEPR2 and the 78 proteins with upregulated phosphorylation

**Supplementary Figure 2.** The transcription levels of various ABA transporters in the CEPR2-OE-9, WT, and *cepr2-1* seedlings, with or without ABA visualized using qRT-PCR.

**Supplementary Figure 3.** The codon-optimized sequence of NRT1.2^loop^ domain.

**Supplementary Figure 4.** The phosphorylation of NRT1.2 did not affect its interaction with CEPR2

**Supplementary Figure 5.** Construction of the complementary materials of *nrt1.2-1*.

**Supplementary Figure 6.** Gene structure of the Group XIV ubiquitin-conjugating enzymes UBC32, UBC33, and UBC34.

**Supplementary Table 1.** The 78 dramatically-upregulated phosphoproteins identified using in phosphorylation mass spectrometry.

**Supplementary Table 2.** The primers used in this study

**Supplementary Excel 1.** The peptides differentially phosphorylated between the CEPR2-OE-9 and WT seedlings identified using mass spectrometry

## References

Ahn, M.Y., Oh, T.R., Seo, D.H., Kim, J.H., Cho, N.H., and Kim, W.T. (2018). Arabidopsis group XIV ubiquitin-conjugating enzymes AtUBC32, AtUBC33, and AtUBC34 play negative roles in drought stress response. J Plant Physiol 230:73–79.

Boursiac, Y., Leran, S., Corratge-Faillie, C., Gojon, A., Krouk, G., and Lacombe, B. (2013). ABA transport and transporters. Trends Plant Sci 18:325–333.

Chen, C., Wu, C., Miao, J., Lei, Y., Zhao, D., Sun, D., Yang, G., Huang, J., and Zheng, C. (2014). Arabidopsis SAG protein containing the MDN1 domain participates in seed germination and seedling development by negatively regulating ABI3 and ABI5. J Exp Bot 65:35–45.

Chen, H., Zou, Y., Shang, Y., Lin, H., Wang, Y., Cai, R., Tang, X., and Zhou, J.M. (2008). Firefly luciferase complementation imaging assay for protein-protein interactions in plants. Plant Physiol 146:368–376.

Chen, H.H., Qu, L., Xu, Z.H., Zhu, J.K., and Xue, H.W. (2018). EL1-like Casein Kinases Suppress ABA Signaling and Responses by Phosphorylating and Destabilizing the ABA Receptors PYR/PYLs in Arabidopsis. Mol Plant 11:706–719.

Cui, F., Liu, L., Li, Q., Yang, C., and Xie, Q. (2012a). UBC32 mediated oxidative tolerance in Arabidopsis. J Genet Genomics 39:415–417.

Cui, F., Liu, L., Zhao, Q., Zhang, Z., Li, Q., Lin, B., Wu, Y., Tang, S., and Xie, Q. (2012b). Arabidopsis ubiquitin conjugase UBC32 is an ERAD component that functions in brassinosteroid-mediated salt stress tolerance. Plant Cell 24:233–244.

Cutler, S.R., Rodriguez, P.L., Finkelstein, R.R., and Abrams, S.R. (2010). Abscisic acid: emergence of a core signaling network. Annu Rev Plant Biol 61:651–679.

Grondin, A., Rodrigues, O., Verdoucq, L., Merlot, S., Leonhardt, N., and Maurel, C. (2015). Aquaporins Contribute to ABA-Triggered Stomatal Closure through OST1-Mediated Phosphorylation. Plant Cell 27:1945–1954.

Huang, N.C., Liu, K.H., Lo, H.J., and Tsay, Y.F. (1999). Cloning and functional characterization of an Arabidopsis nitrate transporter gene that encodes a constitutive component of low-affinity uptake. Plant Cell 11:1381–1392.

Kang, J., Yim, S., Choi, H., Kim, A., Lee, K.P., Lopez-Molina, L., Martinoia, E., and Lee, Y. (2015). Abscisic acid transporters cooperate to control seed germination. Nat Commun 6:8113.

Kanno, Y., Hanada, A., Chiba, Y., Ichikawa, T., Nakazawa, M., Matsui, M., Koshiba, T., Kamiya, Y., and Seo, M. (2012). Identification of an abscisic acid transporter by functional screening using the receptor complex as a sensor. Proc Natl Acad Sci U S A 109:9653–9658.

Kuromori, T., Miyaji, T., Yabuuchi, H., Shimizu, H., Sugimoto, E., Kamiya, A., Moriyama, Y., and Shinozaki, K. (2010). ABC transporter AtABCG25 is involved in abscisic acid transport and responses. Proc Natl Acad Sci U S A 107:2361–2366.

Kuromori, T., Seo, M., and Shinozaki, K. (2018). ABA Transport and Plant Water Stress Responses. Trends Plant Sci 23:513–522.

Li, W., Song, T., Wallrad, L., Kudla, J., Wang, X., and Zhang, W. (2019). Tissue-specific accumulation of pH-sensing phosphatidic acid determines plant stress tolerance. Nat Plants 5:1012–1021.

Obrdlik, P., El-Bakkoury, M., Hamacher, T., Cappellaro, C., Vilarino, C., Fleischer, C., Ellerbrok, H., Kamuzinzi, R., Ledent, V., Blaudez, D., et al. (2004). K+ channel interactions detected by a genetic system optimized for systematic studies of membrane protein interactions. Proc Natl Acad Sci U S A 101:12242–12247.

Pi, E., Qu, L., Hu, J., Huang, Y., Qiu, L., Lu, H., Jiang, B., Liu, C., Peng, T., Zhao, Y., et al. (2016). Mechanisms of Soybean Roots’ Tolerances to Salinity Revealed by Proteomic and Phosphoproteomic Comparisons Between Two Cultivars. Mol Cell Proteomics 15:266–288.

Quintero, F.J., Martinez-Atienza, J., Villalta, I., Jiang, X., Kim, W.Y., Ali, Z., Fujii, H., Mendoza, I., Yun, D.J., Zhu, J.K., et al. (2011). Activation of the plasma membrane Na/H antiporter Salt-Overly-Sensitive 1 (SOS1) by phosphorylation of an auto-inhibitory C-terminal domain. Proc Natl Acad Sci U S A 108:2611–2616.

Raghavendra, A.S., Gonugunta, V.K., Christmann, A., and Grill, E. (2010). ABA perception and signalling. Trends Plant Sci 15:395–401.

Soma, F., Mogami, J., Yoshida, T., Abekura, M., Takahashi, F., Kidokoro, S., Mizoi, J., Shinozaki, K., and Yamaguchi-Shinozaki, K. (2017). ABA-unresponsive SnRK2 protein kinases regulate mRNA decay under osmotic stress in plants. Nat Plants 3:16204.

Teardo, E., Carraretto, L., Moscatiello, R., Cortese, E., Vicario, M., Festa, M., Maso, L., De Bortoli, S., Cali, T., Vothknecht, U.C., et al. (2019). A chloroplast-localized mitochondrial calcium uniporter transduces osmotic stress in Arabidopsis. Nat Plants 5:581–588.

Waadt, R., and Kudla, J. (2008). In Planta Visualization of Protein Interactions Using Bimolecular Fluorescence Complementation (BiFC). CSH Protoc 2008:pdb prot4995.

Wang, P., Zhao, Y., Li, Z., Hsu, C.C., Liu, X., Fu, L., Hou, Y.J., Du, Y., Xie, S., Zhang, C., et al. (2018). Reciprocal Regulation of the TOR Kinase and ABA Receptor Balances Plant Growth and Stress Response. Mol Cell 69:100–112 e106.

Xu, Q., Truong, T.T., Barrero, J.M., Jacobsen, J.V., Hocart, C.H., and Gubler, F. (2016). A role for jasmonates in the release of dormancy by cold stratification in wheat. J Exp Bot 67:3497–3508.

Xu, Q., Yin, S., Ma, Y., Song, M., Song, Y., Mu, S., Li, Y., Liu, X., Ren, Y., Gao, C., et al. (2020). Carbon export from leaves is controlled via ubiquitination and phosphorylation of sucrose transporter SUC2. Proc Natl Acad Sci U S A 117:6223–6230.

Xu, Y., Yu, Z., Zhang, D., Huang, J., Wu, C., Yang, G., Yan, K., Zhang, S., and Zheng, C. (2018). CYSTM, a Novel Non-Secreted Cysteine-Rich Peptide Family, Involved in Environmental Stresses in Arabidopsis thaliana. Plant Cell Physiol 59:423–438.

Yang, G., Yu, Z., Gao, L., and Zheng, C. (2019). SnRK2s at the Crossroads of Growth and Stress Responses. Trends Plant Sci 24:672–676.

Yu, Z., Zhang, D., Xu, Y., Jin, S., Zhang, L., Zhang, S., Yang, G., Huang, J., Yan, K., Wu, C., et al. (2019). CEPR2 phosphorylates and accelerates the degradation of PYR/PYLs in Arabidopsis. J Exp Bot.

Zhang, H., Li, Y., and Zhu, J.K. (2018). Developing naturally stress-resistant crops for a sustainable agriculture. Nat Plants 4:989–996.

Zhang, H., Zhu, H., Pan, Y., Yu, Y., Luan, S., and Li, L. (2014). A DTX/MATE-type transporter facilitates abscisic acid efflux and modulates ABA sensitivity and drought tolerance in Arabidopsis. Mol Plant 7: 1522–1532.

Zheng, L., Chen, Y., Ding, D., Zhou, Y., Ding, L., Wei, J., and Wang, H. (2019). Endoplasmic reticulum-localized UBC34 interaction with lignin repressors MYB221 and MYB156 regulates the transactivity of the transcription factors in Populus tomentosa. BMC Plant Biol 19:97.

