## Supplementary material for "The stability and ABA import activity of NRT1.2 in *Arabidopsis* are regulated by CEPR2 via phosphorylation modification": suplementary data: Suplementary data.pdf

**A**

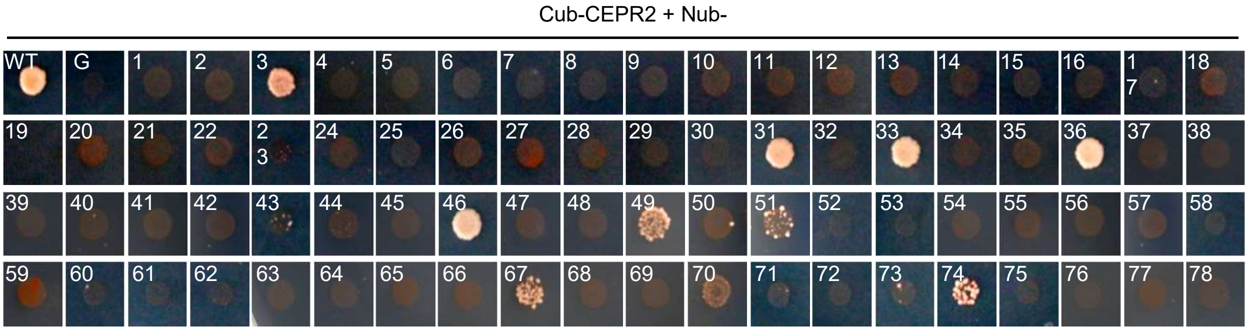


**Supplementary Figure 1. The interaction of 78 proteins with upregulated phosphorylation with CEPR2 by mbSUS**. MbSUS analysis revealed interactions between CEPR2 and 78 target proteins identified in phosphorylation mass spectrometry. Images were taken after four days of culture. The group of “Cub-CEPR2 + NubWT” was used as the positive control. NubG is a mutant form of Nub with weak binding affinity for Cub, and functional ubiquitin can only be reconstituted when NubG and Cub are in close vicinity by fusion with proteins that interact. Thus, the group of “Cub-CEPR2 + NubG” was used as the negative control and further used to identify whether CEPR2 has self-activating activity. 1-78 refers to proteins listed in Supplementary Table 1. MbSUS, Mating-based split ubiquitin system; Cub, the C-termial of ubiquitin; Nub, the N-terminal of ubiquitin.

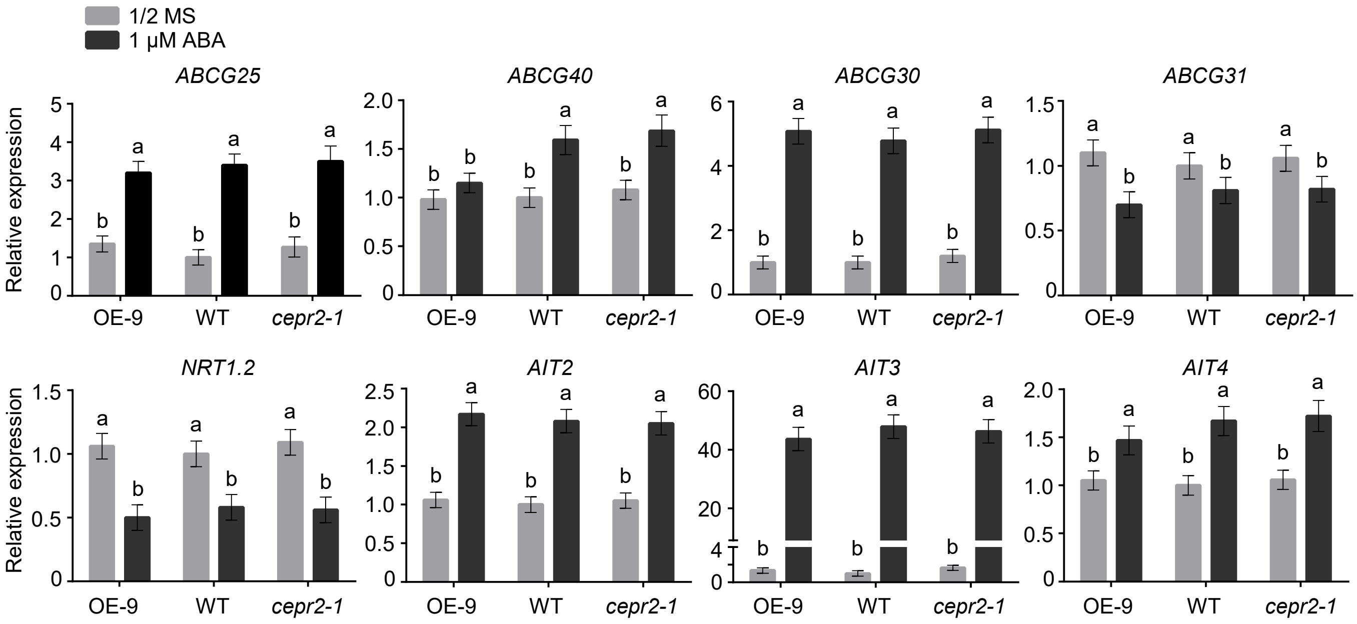


**Supplementary Figure 2. The transcription level of different ABA transporters in CEPR2-OE-9, WT, and *cepr2-1* seedlings with or without ABA were examined by qRT-PCR.**

### (A) The optimized DNA sequence of NRT1.2$^{Loop}$ （Loop domain: 694-1038 bp; 232-346 aa）for antibody preparation

TCTGGATCAAGATTTTATAGGAACAAGATTCCATGTGGAAGTCCTCTCACCACAATCTTGAAGGTTCTTCTTGCGGCTTC

GGTTAAGTGCTGCTCGAGTGGAAGTTCAAGCAATGCGGTTGCGAGTATGTCCGTGAGTCCCTCAAATCATTGCGTATC

AAAGGGGAAAAAAGAAGTTGAATCACAAGGAGAATTGGAAAAGCCACGTCAAGAAGAAGCTTTGCCTCCTCGGGCACA

ACTAACTAACAGTTTGAAAGTATTAAATGGAGCTGCGGATGAAAAACCTGTCCATAGATTGTTAGAATGCACAGTCCAA

CAAGTGGAAGATGTGAAGATTGTCTTGAAA

### (B) Optimized DNA sequence of NRT1.2$^{Loop}$

GGATCCATG TCTGGTTCTCGTTTCTACCGTAACAAAATCCCGTGCGGTTCTCCGCTGACCACCATCCTGAAAGTTCTGCTGGCTGCTT

*Bam*HI

S277

CTGTTAAATGCTGCTCTTCTGGTTCTTCTTCTAACGCTGTTGCTTCTATGTCTGTTTCTCCGTCTAACCACTGCGTTTCTA

S292

AAGGTAAAAAAGAAGTTGAATCTCAGGGTGAACTGGAAAAACCGCGTCAGGAAGAAGCTCTGCCGCCGCGTGCTCAG

CTGACCAACTCTCTGAAAGTTCTGAACGGTGCTGCTGACGAAAAACCGGTTCACCGTCTGCTGGAATGCACCGTTCAG

CAGGTTGAAGACGTTAAAATCGTTCTGAAA TAAGAATTC

*Eco*RI

### (C) Based on the optimized DNA sequence, the mutation of S277A site was accompanied

GGATCCATG TCTGGTTCTCGTTTCTACCGTAACAAAATCCCGTGCGGTTCTCCGCTGACCACCATCCTGAAAGTTCTGCTGGCTGCTT

*Bam*HI

**S277A**

CTGTTAAATGCTGCTCTTCTGGTTCTTCTTCTAACGCTGTTGCTTCTATGTCTGTT**GCT**CCGTCTAACCACTGCGTTTCT

AAAGGTAAAAAAGAAGTTGAATCTCAGGGTGAACTGGAAAAACCGCGTCAGGAAGAAGCTCTGCCGCCGCGTGCTCA

GCTGACCAACTCTCTGAAAGTTCTGAACGGTGCTGCTGACGAAAAACCGGTTCACCGTCTGCTGGAATGCACCGTTCA

GCAGGTTGAAGACGTTAAAATCGTTCTGAAATAAGAATTC

*Eco*RI

### (D) Based on the optimized DNA sequence, the mutation of S292A site was accompanied

GGATCCATG TCTGGTTCTCGTTTCTACCGTAACAAAATCCCGTGCGGTTCTCCGCTGACCACCATCCTGAAAGTTCTGCTGGCTGCTT

*Bam*HI

CTGTTAAATGCTGCTCTTCTGGTTCTTCTTCTAACGCTGTTGCTTCTATGTCTGTTTCTCCGTCTAACCACTGCGTTTCTA

**S292A**

AAGGTAAAAAAGAAGTTGAA**GCT**CAGGGTGAACTGGAAAAACCGCGTCAGGAAGAAGCTCTGCCGCCGCGTGCTCAG

CTGACCAACTCTCTGAAAGTTCTGAACGGTGCTGCTGACGAAAAACCGGTTCACCGTCTGCTGGAATGCACCGTTCAG

CAGGTTGAAGACGTTAAAATCGTTCTGAAA TAAGAATTC

*Eco*RI

**Supplementary Figure 3. The optimizing codon sequence of loop domain of NRT1.2**

**(A)** The codon sequence of cytoplasmic domain of NRT1.2 downloaded from Tair 10 website.

**(B)** The optimized DNA sequence of cytoplasmic domain of NRT1.2 (http://www.jcat.de/Start.jsp).

**(C)** Candidate phosphorylation site (S277) is mutated to alanine (S277A), which is highlighted in red in the optimized DNA sequence of

cytoplasmic domain of NRT1.2.

**(D)** Candidate phosphorylation site (S292) is mutated to alanine (S292A), which is highlighted in red in the optimized DNA sequence of

cytoplasmic domain of NRT1.2.

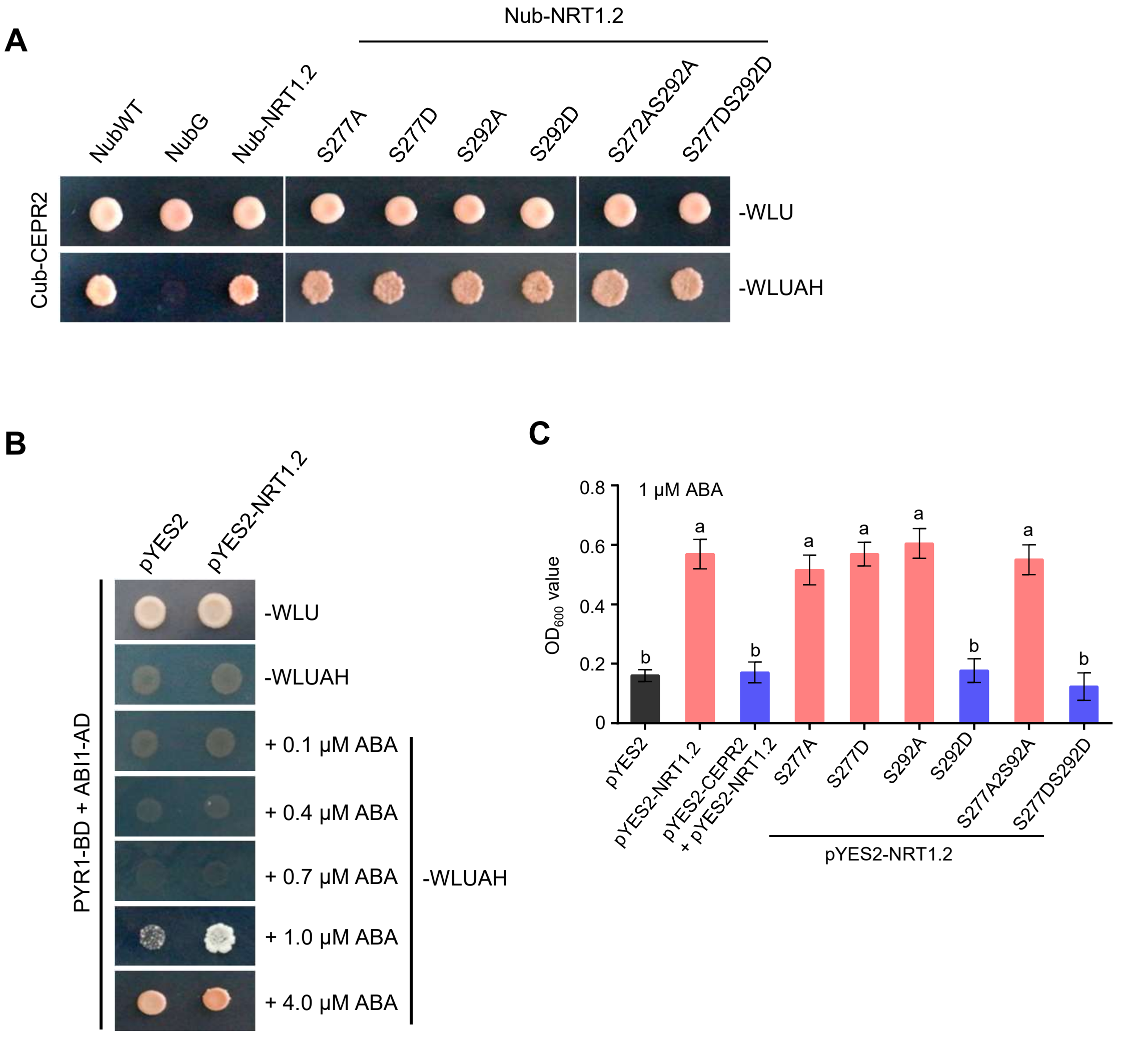


**Supplementary Figure 4. The phosphorylation of NRT1.2 did not affect the interaction with CEPR2**

**(A)** MbSUS assay shows the interaction ability between CEPR2 and phosphomimic mutated forms of NRT1.2. Yeast transformants were spotted on the control medium SD-Trp/-Leu/-Ura (SD-WLU) and selection medium SD-Trp/-Leu/-Ura/-Ade/-His (SD-WLUAH). Images were taken after culturing at 30°C for four days.

**(B)** Yeast transformants were spotted on the control medium SD-WLU and selection medium SD-WLUAH with indicated concentrations of ABA. Images were taken after culturing at 30°C for four days.

**(C)** Optical densities (OD) at 600 nm of the independent transformants yeast cells in Figure5a. Error bars indicate SEM (N = 3). Bars labeled with different lowercase letters are significantly different from one another ($P < 0.05$; one way ANOVA).

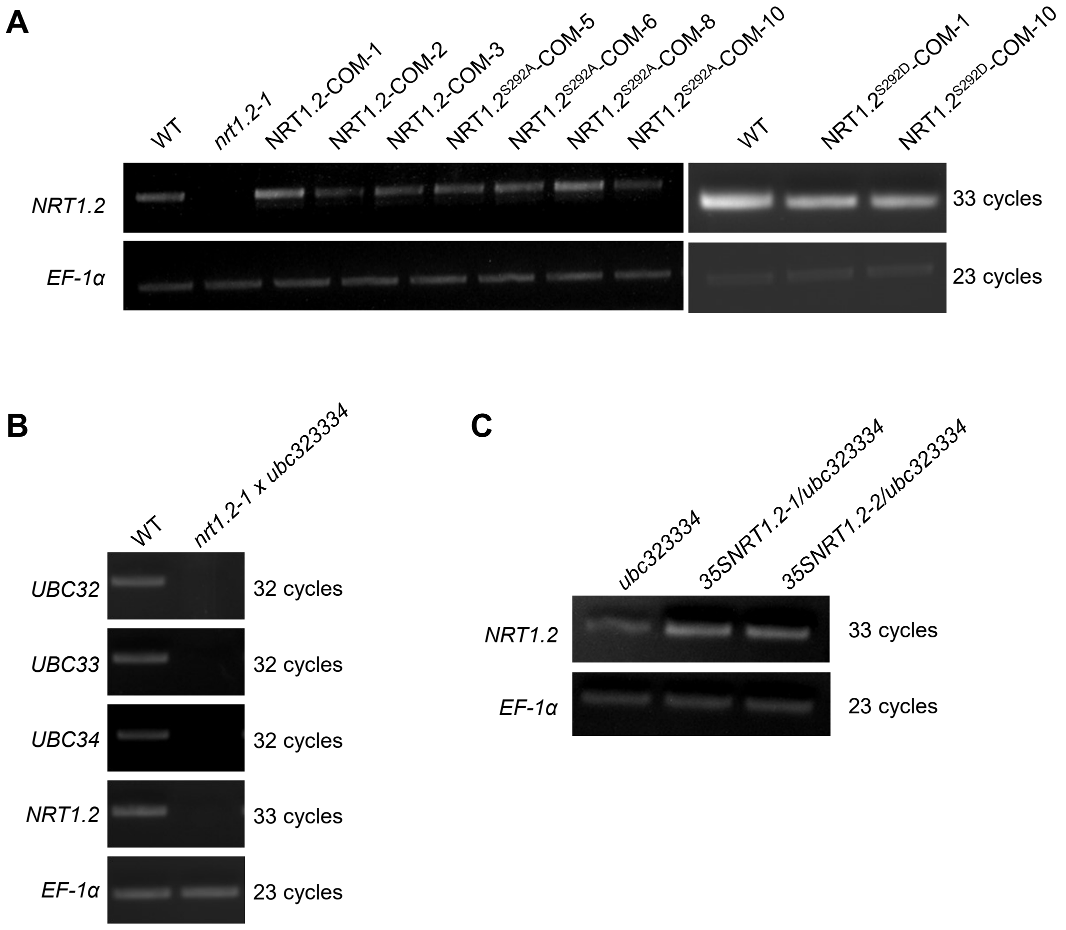


**Supplementary Figure 5. Construction of the complementary materials of *nrt1.2-1***

**(A)** The expression levels of *NRT1.2* in different complementary materials carrying native promoter-driven native NRT1.2 ,phosphomimic mutant and nonphosphomimic mutant of *NRT1.2* were detected by RT-PCR. *EF-1α* was used as the internal control.

**(B)** The expression levels of *UBC32, UBC33, UBC34* and *NRT1.2* in *nrt1.2-1ubc323334* quardruple mutants were detected by RT-PCR. *EF-1α* was used as the internal control.

**(C)** The expression levels of *NRT1.2* in *35SNRT1.2/ubc323334* mutants were detected by RT-PCR. *EF-1α* was used as the internal control.

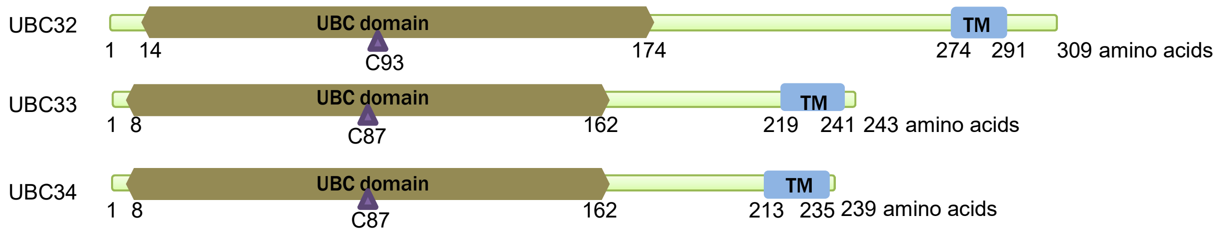


**Supplementary Figure 6. Gene structure of Group XIV ubiquitin-conjugating enzymes UBC32, UBC33 and UBC34.** The N-terminal UBC domain and C-terminal TM motif are indicated by UBC32, UBC33 and UBC34 whose UBC domain include a conserved cysteine ( C93 or C87 ) that can carry the ubiquitins.

**Supplementary Table 1. The 78 dramatically upregulated phosphoproteins identified in phosphorylation mass spectrometry**

| Protein number | Gene ID | Notes on the TAIR website |
|---|---|---|
| 1 | AT3G63400 | Cyclophilin-like peptidyl-prolyl cis-trans isomerase family protein |
| 2 | AT1G35580 | CINV1, carbohydrate metabolic process |
| 3 | AT1G70770 | Unknown function |
| 4 | AT4G38470 | SERINE/THREONINE/TYROSINE KINASE 46, STY46 |
| 5 | AT4G10120 | ATSPS4F, SUCROSE PHOSPHATE SYNTHASE 4F |
| 6 | AT3G19820 | DWF1, involved in brassinosteroid biosynthetic process. |
| 7 | AT4G02510 | PPI2, involved in protein import into chloroplast stroma |
| 8 | AT1G37130 | NR2, has nitrate reductase activity |
| 9 | AT3G17160 | Unknown function |
| 10 | AT3G44750 | HD2A, histone deacetylase |
| 11 | AT5G47430 | DWNN domain, a CCHC-type zinc finger |
| 12 | AT5G22650 | HD2B, encodes a histone deacetylase |
| 13 | AT1G31830 | ATPUT2, encodes POLYAMINE UPTAKE TRANSPORTER 2 |
| 14 | AT1G13320 | Protein phosphatase 2A subunit A3 |
| 15 | AT5G64200 | ATSC35, ORTHOLOG OF HUMAN SPLICING FACTOR SC35 |
| 16 | AT4G13510 | AMT1;1, AMMONIUM TRANSPORT 1 |
| 17 | AT2G31890 | RAP, functions as a negative regulator of plant disease resistance |
| 18 | AT4G22670 | HIP1, HSP70-interacting protein 1 |
| 19 | AT2G19490 | recA DNA recombination family protein |
| 20 | AT4G25890 | Invoved in translational elongation |
| 21 | AT1G01100 | 60S acidic ribosomal protein P1-1 |
| 22 | AT5G08610 | PDE340, helicase activity |
| 23 | AT1G15440 | PWP2, ribosome biogenesis co-factor |
| 24 | AT1G68830 | STN7, regulation of photosynthesis |
| 25 | AT5G09850 | Transcription elongation factor family |
| 26 | AT4G13200 | Unknown protein |
| 27 | AT4G26780 | Response to heat, unknown function |
| 28 | AT1G56220 | Dormancy associated family protein |
| 29 | AT3G50590 | WD40/YVTN repeat protein; involved in pollen development |
| 30 | AT1G19870 | IQD32, microtubule-associated protein |
| 31 | AT5G14120 | Unknown protein |
| 32 | AT3G56150 | eIF3c - eukaryotic initiation factor 3c |
| 33 | AT2G16850 | PIP2;8, Aquaporins, Water transporter |
| 34 | AT3G61050 | ATCLB, negative regulation of transcription |
| 35 | AT1G12920 | Eukaryotic release factor 1-2 |
| 36 | AT3G17650 | YELLOW STRIPE LIKE 5, YSL5 metal-nicotianamine transporter |
| 37 | AT2G27710 | 60S acidic ribosomal protein family |
| 38 | AT3G25500 | AFH1, actin cytoskeleton organization |
| 39 | AT2G37340 | RSZ33, spliceosomal complex assembly |

| Protein number | Gene ID | Notes on the TAIR website |
|---|---|---|
| 40 | AT5G23060 | CAS, regulation of stomatal closure |
| 41 | AT3G08940 | LHCB4.2, involved in photosynthesis |
| 42 | AT4G39680 | SAP domain-containing protein |
| 43 | AT3G51800 | ERBB-3 BINDING PROTEIN 1 |
| 44 | AT4G37190 | Unknown function |
| 45 | AT1G43690 | Lys48-specific deubiquitinase activity |
| 46 | AT1G69850 | NRT1.2, ABA and $NO_3^-$ transporter |
| 47 | AT5G19500 | Involved in amino acid transport |
| 48 | AT1G22530 | PATL2, response to auxin stimulus |
| 49 | AT2G39010 | PIP2;6, Aquaporins, Water transporter |
| 50 | AT1G80270 | PENTATRICOPEPTIDE REPEAT 596 |
| 51 | AT5G12080 | ATMSL10, Mechanosensitive Ion Channel |
| 52 | AT3G49601 | Pre-mRNA-splicing factor |
| 53 | AT3G12980 | HAC5, histone acetylation |
| 54 | AT3G13570 | SC35-LIKE SPLICING FACTOR 30A |
| 55 | AT4G31420 | REIL1, positive regulation of ribosome biogenesis |
| 56 | AT4G21660 | Proline-rich spliceosome-associated family protein |
| 57 | AT2G26460 | SMU2, involved in RNA splicing. |
| 58 | AT1G76850 | EXOCYST COMPLEX COMPONENT SEC5, SEC5A |
| 59 | AT1G06210 | ENTH/VHS/GAT family protein |
| 60 | AT1G20220 | Alba DNA/RNA-binding protein |
| 61 | AT3G57150 | NAP57, a putative pseudouridine synthase |
| 62 | AT2G39810 | HOS1, Functions as an E3 ligase required for the ubiquitination of ICE1 |
| 63 | AT3G53500 | RSZ32, Splicing Factors (SR proteins) |
| 64 | AT1G59710 | Unknown function |
| 65 | AT4G29190 | OXIDATION-RELATED ZINC FINGER 2 |
| 66 | AT1G48920 | NUCLEOLIN LIKE 1 |
| 67 | AT3G21770 | Peroxidase superfamily protein |
| 68 | AT1G16610 | SR45, a spliceosome protein |
| 69 | AT1G13650 | Unknown function |
| 70 | AT1G45230 | DCL protein, chloroplast rRNA processing |
| 71 | AT5G56000 | ATHSP90.4, response to heat |
| 72 | AT1G03380 | ATG18G, autophagy, protein transport |
| 73 | AT5G62190 | ATRH7, RNA metabolic process |
| 74 | AT3G55770 | WLIM2B, Regulates actin cytoskeleton organization. |
| 75 | AT3G09200 | Ribosomal protein L10 family protein, involved in cytoplasmic translation |
| 76 | AT1G77760 | NR1, has nitrate reductase activity |
| 77 | AT1G72160 | Sec14p-like phosphatidylinositol transfer family protein |
| 78 | AT3G17205 | Ubiquitin protein ligase 6 |

**Supplementary Table 2. The primers used in this study**

| Purpose | Name | Sequence (5'-3') |
|---|---|---|
| qRT ABA transporters | ABCG25 F | GTGGTTACTACGTCAACA |
| | ABCG25 R | CCTTGCTTACCCTTTGAA |
| | ABCG30 F | GAGTTCGCAATATGGAGATG |
| | ABCG30 R | CACAACAGCCAATGATTCA |
| | ABCG31 F | GGTATGTATGCTCCAATTCC |
| | ABCG31 R | CAATGGTGAAGTATGTGATGA |
| | ABCG40 F | CACCTTCTACGGAATGAT |
| | ABCG40 R | CAAAGCCAGTAGTACCAT |
| | NRT1.2 F | CACCTTCCTCAATGAGAT |
| | NRT1.2 R | ACATTAGCCAATAGAAGTAGT |
| | AIT2 F | TGATCTGTTCACACTAGC |
| | AIT2 R | CTTTGTCCAGCACTCTTA |
| | AIT3 F | TTAGCTCTCACCTCTATTTC |
| | AIT3 R | AGATTCTCTAAAGAAGAACTCTA |
| | AIT4 F | ACCTTTATACGAATTCTTTGG |
| | AIT4 R | GAGATTCTGAAGTTGTTATGAA |
| | UBQ10 F | GCCAAGATCCAGGACAAGG |
| | UBQ10 R | CGCAGGACCAAGTGAAGAG |
| | UBC21 F | CTGCGACTCAGGGAATCTTCTAA |
| | UBC21 R | TTGTGCCATTGAATTGAACCC |
| MbSUS ABA transporters | Nub-ABCG25 F | acaagtttgtacaaaaaagcaggctctccaaccaccATGTCAGCTTTTGACGGCG |
| | Nub-ABCG25 R | tccgccaccaccaaccactttgtacaagaaagctgggtaTTAATGTTTGATACGTCTCAAAGCTAG |
| | Nub-ABCG30 F | acaagtttgtacaaaaaagcaggctctccaaccaccATGATCCAAACAGGTGAAGAAGATG |
| | Nub-ABCG30 R | tccgccaccaccaaccactttgtacaagaaagctgggtaTTTCTTTTGGAAACTGAGTTTGCTCA |
| | Nub-ABCG31 F | acaagtttgtacaaaaaagcaggctctccaaccaccATGGCGGCGGCTTCGAAT |
| | Nub-ABCG31 R | tccgccaccaccaaccactttgtacaagaaagctgggtaTCTTCTCTGGAAGTTGAGGTATTTGAC |
| | Nub-ABCG40 F | acaagtttgtacaaaaaagcaggctctccaaccaccATGGAGGGAACTAGTTTTCACC |
| | Nub-ABCG40 R | tccgccaccaccaaccactttgtacaagaaagctgggtaCTATCGTTTTTGGAAATTGAAACTCTTG |
| | Nub-NRT1.2 F | acaagtttgtacaaaaaagcaggctctccaaccaccATGGAAGTGGAAGAAGAGG |
| | Nub-NRT1.2 R | tccgccaccaccaaccactttgtacaagaaagctgggtaTTAGCTTCTTGAACCAGTTG |
| | Nub-AIT2 F | acaagtttgtacaaaaaagcaggctctccaaccaccATGGAAGTAGAAATGCATGGTG |
| | Nub-AIT2 R | tccgccaccaccaaccactttgtacaagaaagctgggtaTCAACTTATTGAACCAGTTGAGATATAC |
| | Nub-AIT3 F | acaagtttgtacaaaaaagcaggctctccaaccaccATGCAGATTGAGATGGAAGA |
| | Nub-AIT3 R | tccgccaccaccaaccactttgtacaagaaagctgggtaCTAATATCTTTTCGCCCAGA |
| | Nub-AIT4 F | acaagtttgtacaaaaaagcaggctctccaaccaccATGGAGAATGATATGGAAGAGAA |
| | Nub-AIT4 R | tccgccaccaccaaccactttgtacaagaaagctgggtaTTAATATCTCTTTGCCCAAAAAATATAG |
| | Cub-CEPR2 F | acaagtttgtacaaaaaagcaggctctccaaccaccATGTCGAGAAGACCAGACCT |
| | Cub-CEPR2 R | tccgccaccaccaaccactttgtacaagaaagctgggtaTACTGTAATCTTTCCAGTTGTGTC |
| RT-PCR T-DNA lines | *nrt1.2* F | TTTTCTCCTAGCTCTCCTCGG |
| | *nrt1.2* R | TGACAAGGAGATTATAAGTTAAGTCAGC |
| | ubc32 F | GTATGGCTCTCCTGAACGGC |
| | ubc32 R | CGATGGAAGGTGCTTGGCTA |
| | ubc33 F | GTGTAGCCGAGAAACAACGG |
| | ubc33 R | TGTTGAGTGGCTGCTTCTTCT |
| | ubc34 F | TCCCATGTTGTTGCTCGTCC |
| | ubc34 R | GGTGTTCCCTCACTGCCTTC |
| | EF-1α F | gtatggttgttacctttgctcccacag |
| | EF-1α R | catcatttggcacccttcttcactgc |
| MbSUS<br><br>78 Proteins identified in phos-mass spectrometry | Nub-1 F | acaagtttgtacaaaaaagcaggctctccaaccaccATGACTAAAAAGAAGAATCCTAA |
| | Nub-1 R | tccgccaccaccaaccactttgtacaagaaagctgggtaATCCGCATAGCTAACCAGA |
| | Nub-2 F | acaagtttgtacaaaaaagcaggctctccaaccaccATGGAAGGTGTTGGACTAAGAGC |
| | Nub-2 R | tccgccaccaccaaccactttgtacaagaaagctgggtaGAGTTGTGGCCAAGACGCA |
| | Nub-3 F | acaagtttgtacaaaaaagcaggctctccaaccaccATGGATCCGATCGAATCTGTCG |
| | Nub-3 R | tccgccaccaccaaccactttgtacaagaaagctgggtaCTTCTTCAAAGCTGTGGAGA |
| | Nub-4 F | acaagtttgtacaaaaaagcaggctctccaaccaccATGGTGATGGAGGACAACG |
| | Nub-4 R | tccgccaccaccaaccactttgtacaagaaagctgggtaATGATGTGTGGTGCTTCTCC |
| | Nub-5 F | acaagtttgtacaaaaaagcaggctctccaaccaccATGGCAAGAAATGATTGGATAAA |
| | Nub-5 R | tccgccaccaccaaccactttgtacaagaaagctgggtaCTTGATCCCATAGGCCTCT |
| | Nub-6 F | acaagtttgtacaaaaaagcaggctctccaaccaccATGTCGGATCTTCAGACACCG |
| | Nub-6 R | tccgccaccaccaaccactttgtacaagaaagctgggtaATCTGCCTCGGCATAAGCAG |
| | Nub-7 F | acaagtttgtacaaaaaagcaggctctccaaccaccATGGACTCAAAGTCGGTTA |
| | Nub-7 R | tccgccaccaccaaccactttgtacaagaaagctgggtaGTACATGCTGTACTTGTCGT |
| | Nub-8 F | acaagtttgtacaaaaaagcaggctctccaaccaccATGGCGGCCTCTGTAGAT |
| | Nub-8 R | tccgccaccaccaaccactttgtacaagaaagctgggtaGAATATCAAGAAATCCTCCT |
| | Nub-9 F | acaagtttgtacaaaaaagcaggctctccaaccaccATGACGCATGTTGAAGATGATA |
| | Nub-9 R | tccgccaccaccaaccactttgtacaagaaagctgggtaCCGTTTAGAAGGCCTAATGT |

⋯ continued ⋯

| Purpose | Name | Sequence (5'-3') |
|---|---|---|
| MbSUS 78 Proteins identified in phos-mass spectrometry | 10 F | acaagtttgtacaaaaaagcaggctctccaaccaccATGGAGTTCTGGGGAATTGAAGT |
| | 10 R | tccgccaccaccaaccactttgtacaagaaagctgggtaCTTGGCAGCAGCGTGCTTG |
| | 11 F | acaagtttgtacaaaaaagcaggctctccaaccaccATGGCAATTTATTACAAGTTTAA |
| | 11 R | tccgccaccaccaaccactttgtacaagaaagctgggtaAGCTCGAGATCTCTCCC |
| | 12 F | acaagtttgtacaaaaaagcaggctctccaaccaccATGGAGTTCTGGGGAGTTGC |
| | 12 R | tccgccaccaccaaccactttgtacaagaaagctgggtaAGCTCTACCCTTTCCCTTGC |
| | 13 F | acaagtttgtacaaaaaagcaggctctccaaccaccATGCAGAAGCGGAGAATC |
| | 13 R | tccgccaccaccaaccactttgtacaagaaagctgggtaACGTATTAGAGTTTCTTCGT |
| | 14 F | acaagtttgtacaaaaaagcaggctctccaaccaccATGTCTATGGTTGATGAGC |
| | 14 R | tccgccaccaccaaccactttgtacaagaaagctgggtaGCTAGACATCATCACATTGT |
| | 15 F | acaagtttgtacaaaaaagcaggctctccaaccaccATGTCGCACTTCGGAAGGTC |
| | 15 R | tccgccaccaccaaccactttgtacaagaaagctgggtaTTCCGCAGCATAAGGAGA |
| | 16 F | acaagtttgtacaaaaaagcaggctctccaaccaccATGTCTTGCTCGGCCAC |
| | 16 R | tccgccaccaccaaccactttgtacaagaaagctgggtaAACCGGAGTAGGTGTAGTAT |
| | 17 F | acaagtttgtacaaaaaagcaggctctccaaccaccATGGAGTGTGTAGTTCCA |
| | 17 R | tccgccaccaccaaccactttgtacaagaaagctgggtaTATGCAGCCGGTGAGAA |
| | 18 F | acaagtttgtacaaaaaagcaggctctccaaccaccATGGATTCAACGAAGCTTAG |
| | 18 R | tccgccaccaccaaccactttgtacaagaaagctgggtaCTGAGGTCCTGCAAATTT |
| | 19 F | acaagtttgtacaaaaaagcaggctctccaaccaccATGGCGAGGATTCTCCGAA |
| | 19 R | tccgccaccaccaaccactttgtacaagaaagctgggtaTGCTGCCTCAACAACCACT |
| | 20 F | acaagtttgtacaaaaaagcaggctctccaaccaccATGGGAGTATTCACATTCGTA |
| | 20 R | tccgccaccaccaaccactttgtacaagaaagctgggtaACCAAAGAGATCGAATCCG |
| | 21 F | acaagtttgtacaaaaaagcaggctctccaaccaccATGTCGACAGTTGGAGAGC |
| | 21 R | tccgccaccaccaaccactttgtacaagaaagctgggtaGTCAAACAAACCGAAACCC |
| | 22 F | acaagtttgtacaaaaaagcaggctctccaaccaccATGTCCTCGAAGTTCCCT |
| | 22 R | tccgccaccaccaaccactttgtacaagaaagctgggtaCTTGGTTCTAAGACCAGGA |
| | 23 F | acaagtttgtacaaaaaagcaggctctccaaccaccATGGAGTTCCGTTTCGAG |
| | 23 R | tccgccaccaccaaccactttgtacaagaaagctgggtaATGATTGTTTGGAACAGAAC |
| | 24 F | acaagtttgtacaaaaaagcaggctctccaaccaccATGGCTACAATATCTCCGGG |
| | 24 R | tccgccaccaccaaccactttgtacaagaaagctgggtaCTCCTCTCTGGGGATCCA |
| | 25 F | acaagtttgtacaaaaaagcaggctctccaaccaccATGGATTTGGATGATTTCCGAT |
| | 25 R | tccgccaccaccaaccactttgtacaagaaagctgggtaCCAGTGTCTACCACCTTGA |
| | 26 F | acaagtttgtacaaaaaagcaggctctccaaccaccATGAGTAGCTTCACGATTCCAT |
| | 26 R | tccgccaccaccaaccactttgtacaagaaagctgggtaGTCTTCATCACTGGAACTCG |
| | 27 F | acaagtttgtacaaaaaagcaggctctccaaccaccATGTTGGTTTTGAGAATTTTGT |
| | 27 R | tccgccaccaccaaccactttgtacaagaaagctgggtaAGCATCAGACTCTTTCTTTT |
| | 28 F | acaagtttgtacaaaaaagcaggctctccaaccaccATGAATCGAAGAACCTCTACA |
| | 28 R | tccgccaccaccaaccactttgtacaagaaagctgggtaCATGCCGTAAGTAGGAGG |
| | 29 F | acaagtttgtacaaaaaagcaggctctccaaccaccATGGAGTGGGCAACGGTG |
| | 29 R | tccgccaccaccaaccactttgtacaagaaagctgggtaGCCAAACGGTGTGGATAC |
| | 30 F | acaagtttgtacaaaaaagcaggctctccaaccaccATGGGAAGATCTCCAGCT |
| | 30 R | tccgccaccaccaaccactttgtacaagaaagctgggtaCCTCTGCCATTTTCTATCC |
| | 31 F | acaagtttgtacaaaaaagcaggctctccaaccaccATGGCGTCGACTACACGAG |
| | 31 R | tccgccaccaccaaccactttgtacaagaaagctgggtaGGTGCGGGTCTTGCCATA |
| | 32 F | acaagtttgtacaaaaaagcaggctctccaaccaccATGACGTCTCGTTTTTTCACT |
| | 32 R | tccgccaccaccaaccactttgtacaagaaagctgggtaAGTACGAACACCTCGGTT |
| | 33 F | acaagtttgtacaaaaaagcaggctctccaaccaccATGTCAAAAGAAGTGAGTGAAG |
| | 33 R | tccgccaccaccaaccactttgtacaagaaagctgggtaATTGGTTGGGTTGCTGCG |
| | 34 F | acaagtttgtacaaaaaagcaggctctccaaccaccATGGGTTTGATTTCTGGGATTC |
| | 34 R | tccgccaccaccaaccactttgtacaagaaagctgggtaCTGCTGTTTTGCACCATCG |
| | 35 F | acaagtttgtacaaaaaagcaggctctccaaccaccATGGCAGAAGAAGCGGAT |
| | 35 R | tccgccaccaccaaccactttgtacaagaaagctgggtaATCAGAATCTTCGTAAACTT |
| | 36 F | acaagtttgtacaaaaaagcaggctctccaaccaccATGAGAAAGGGAGTTCTAAATC |
| | 36 R | tccgccaccaccaaccactttgtacaagaaagctgggtaAATGGATCCTTTCAGGAAG |
| | 37 F | acaagtttgtacaaaaaagcaggctctccaaccaccATGAAGGTTGTTGCCGCA |
| | 37 R | tccgccaccaccaaccactttgtacaagaaagctgggtaCTCGAATAGACTGAAACCC |
| | 38 F | acaagtttgtacaaaaaagcaggctctccaaccaccATGCTCTTCTTCTTATTCTT |
| | 38 R | tccgccaccaccaaccactttgtacaagaaagctgggtaAGAAACTAATGAGATTGAGT |
| | 39 F | acaagtttgtacaaaaaagcaggctctccaaccaccATGCCTCGCTATGATGATCG |
| | 39 R | tccgccaccaccaaccactttgtacaagaaagctgggtaAGGAGACTCACTTCCTCT |
| | 40 F | acaagtttgtacaaaaaagcaggctctccaaccaccATGGCTATGGCGGAAATGG |
| | 40 R | tccgccaccaccaaccactttgtacaagaaagctgggtaGTCGGAGCTAGGAAGGAAC |
| | 41 F | acaagtttgtacaaaaaagcaggctctccaaccaccATGGCCGCCACTTCAACC |
| | 41 R | tccgccaccaccaaccactttgtacaagaaagctgggtaGGAGGAAGAGAAGGTATCGA |
| | 42 F | acaagtttgtacaaaaaagcaggctctccaaccaccATGTCGTCATCGCCTTTT |
| | 42 R | tccgccaccaccaaccactttgtacaagaaagctgggtaCTTGTTATTATTCGCTGCAA |
| | 43 F | acaagtttgtacaaaaaagcaggctctccaaccaccATGAGTTCGGACGATGAG |
| | 43 R | tccgccaccaccaaccactttgtacaagaaagctgggtaTTCTTGAGCATTACTACTTG |

··· continued ···

| Purpose | Name | Sequence (5'-3') |
|---|---|---|
| | 44 F | acaagtttgtacaaaaaagcaggctctccaaccaccATGAGAGAAATTGTGACGAT |
| | 44 R | tccgccaccaccaaccactttgtacaagaaagctgggtaATCTGAATCCGAAGATTCGA |
| | 45 F | acaagtttgtacaaaaaagcaggctctccaaccaccATGGCGGATCATCAAGAA |
| | 45 R | tccgccaccaccaaccactttgtacaagaaagctgggtaGACTATACTTGGAGGATCTC |
| | 46 F | acaagtttgtacaaaaaagcaggctctccaaccaccATGGAAGTGGAAGAAGAGG |
| | 46 R | tccgccaccaccaaccactttgtacaagaaagctgggtaTTAGCTTCTTGAACCAGTTG |
| | 47 F | acaagtttgtacaaaaaagcaggctctccaaccaccATGTCTGTGTCTCTAAGC |
| | 47 R | tccgccaccaccaaccactttgtacaagaaagctgggtaAGAAACATTTAGATACTTTG |
| | 48 F | acaagtttgtacaaaaaagcaggctctccaaccaccATGGCTCAAGAAGAGATACAGA |
| | 48 R | tccgccaccaccaaccactttgtacaagaaagctgggtaTGCTTGGGTTTTGGACCT |
| | 49 F | acaagtttgtacaaaaaagcaggctctccaaccaccATGACGAAGGATGAGTTGACG |
| | 49 R | tccgccaccaccaaccactttgtacaagaaagctgggtaAGCATGGAGCTCATGAAGCT |
| | 50 F | acaagtttgtacaaaaaagcaggctctccaaccaccATGTTCGCTCTTTCCAAG |
| | 50 R | tccgccaccaccaaccactttgtacaagaaagctgggtaATCCAGAATATCAGAGATAG |
| | 51 F | acaagtttgtacaaaaaagcaggctctccaaccaccATGGCAGAACAAAAGAGT |
| | 51 R | tccgccaccaccaaccactttgtacaagaaagctgggtaGTTCTTCTTTGTGAGATTAA |
| | 52 F | acaagtttgtacaaaaaagcaggctctccaaccaccATGTACAACGGAATAGGGT |
| | 52 R | tccgccaccaccaaccactttgtacaagaaagctgggtaACCATGCGAGGATCTCTT |
| | 53 F | acaagtttgtacaaaaaagcaggctctccaaccaccATGGCTCAGGGGCAGAATAGG |
| | 53 R | tccgccaccaccaaccactttgtacaagaaagctgggtaTTCAGGAGTGGAGGCCGT |
| | 54 F | acaagtttgtacaaaaaagcaggctctccaaccaccATGAGAGGAAGGAGCTACACG |
| | 54 R | tccgccaccaccaaccactttgtacaagaaagctgggtaCTGGCTTGGAGAACGGTC |
| | 55 F | acaagtttgtacaaaaaagcaggctctccaaccaccATGCCTGGTTTAACATGTAAC |
| | 55 R | tccgccaccaccaaccactttgtacaagaaagctgggtaATATGGGACGTTATTGGGC |
| | 56 F | acaagtttgtacaaaaaagcaggctctccaaccaccATGACCGCCGATTCAAC |
| | 56 R | tccgccaccaccaaccactttgtacaagaaagctgggtaGGAAGAAGAAGAAAGATTTC |
| | 57 F | acaagtttgtacaaaaaagcaggctctccaaccaccATGAAACCTTCAAAATCGCAT |
| | 57 R | tccgccaccaccaaccactttgtacaagaaagctgggtaATGCTTGGATCTCTTAGGAG |
| | 58 F | acaagtttgtacaaaaaagcaggctctccaaccaccATGTCGAGCGATAGCAATGATCT |
| | 58 R | tccgccaccaccaaccactttgtacaagaaagctgggtaTCTTCGTCTGGGTCGGGC |
| | 59 F | acaagtttgtacaaaaaagcaggctctccaaccaccATGGACAAATTGAAGATAGCAG |
| | 59 R | tccgccaccaccaaccactttgtacaagaaagctgggtaTTTTTCATCTTCATCACTCG |
| MbSUS | 60 F | acaagtttgtacaaaaaagcaggctctccaaccaccATGGATAAGTATCAGAGAGTTGA |
| | 60 R | tccgccaccaccaaccactttgtacaagaaagctgggtaTGCAGCAGCCTGGATAGG |
| 78 Proteins identified in phos-mass spectrometry | 61 F | acaagtttgtacaaaaaagcaggctctccaaccaccATGGCGGAGGTCGACATC |
| | 61 R | tccgccaccaccaaccactttgtacaagaaagctgggtaTTCCTCATCATCCTCACTG |
| | 62 F | acaagtttgtacaaaaaagcaggctctccaaccaccATGGATACGAGAGAAATCAACG |
| | 62 R | tccgccaccaccaaccactttgtacaagaaagctgggtaTCTTGCTGCGAATCTACGT |
| | 63 F | acaagtttgtacaaaaaagcaggctctccaaccaccATGCCTCGCTATGATGATCG |
| | 63 R | tccgccaccaccaaccactttgtacaagaaagctgggtaAGGTGACTCACTGCCTTT |
| | 64 F | acaagtttgtacaaaaaagcaggctctccaaccaccATGGAGATTTTTCAAAAGGC |
| | 64 R | tccgccaccaccaaccactttgtacaagaaagctgggtaCTTGATGAACTTTTCTGCTA |
| | 65 F | acaagtttgtacaaaaaagcaggctctccaaccaccATGATGATCGGAGAAACTCGC |
| | 65 R | tccgccaccaccaaccactttgtacaagaaagctgggtaCATGAGCAGGTCAGATACCC |
| | 66 F | acaagtttgtacaaaaaagcaggctctccaaccaccATGGGAAAGTCTAAATCCGC |
| | 66 R | tccgccaccaccaaccactttgtacaagaaagctgggtaCTCGTCACCGAAGGTAGT |
| | 67 F | acaagtttgtacaaaaaagcaggctctccaaccaccATGAAGACGATGACGCAATTAAA |
| | 67 R | tccgccaccaccaaccactttgtacaagaaagctgggtaACTTCCAGCGACAGAACACC |
| | 68 F | acaagtttgtacaaaaaagcaggctctccaaccaccATGGCGAAACCAAGTCGT |
| | 68 R | tccgccaccaccaaccactttgtacaagaaagctgggtaAGTTTTACGAGGTGGAGG |
| | 69 F | acaagtttgtacaaaaaagcaggctctccaaccaccATGCCTGCATACATTTCT |
| | 69 R | tccgccaccaccaaccactttgtacaagaaagctgggtaGAATCTACAACAGAGAAGAG |
| | 70 F | acaagtttgtacaaaaaagcaggctctccaaccaccATGAGCTTGGCTTCGAT |
| | 70 R | tccgccaccaccaaccactttgtacaagaaagctgggtaTCTGTTCTGCCTACGTTT |
| | 71 F | acaagtttgtacaaaaaagcaggctctccaaccaccATGGCGGACGCAGAGACC |
| | 71 R | tccgccaccaccaaccactttgtacaagaaagctgggtaGTCGACTTCCTCCATCTTGC |
| | 72 F | acaagtttgtacaaaaaagcaggctctccaaccaccATGATGAAGAAGGGGAAAGG |
| | 72 R | tccgccaccaccaaccactttgtacaagaaagctgggtaATCACCTACAAAGGAAACCA |
| | 73 F | acaagtttgtacaaaaaagcaggctctccaaccaccATGCCTTCCCTAATGTTATCTG |
| | 73 R | tccgccaccaccaaccactttgtacaagaaagctgggtaATATCTCTGGCCTCTACCAC |
| | 74 F | acaagtttgtacaaaaaagcaggctctccaaccaccATGTCTTTTACAGGAACTCAAC |
| | 74 R | tccgccaccaccaaccactttgtacaagaaagctgggtaAGATTCAGGAACGGAGGC |
| | 75 F | acaagtttgtacaaaaaagcaggctctccaaccaccATGGTGAAAGCAACTAAGGC |
| | 75 R | tccgccaccaccaaccactttgtacaagaaagctgggtaCTCTTCATCGAACAAACCG |
| | 76 F | acaagtttgtacaaaaaagcaggctctccaaccaccATGGCGACCTCCGTCGAT |
| | 76 R | tccgccaccaccaaccactttgtacaagaaagctgggtaGAAGATTAAGAGATCCTCCT |
| | 77 F | acaagtttgtacaaaaaagcaggctctccaaccaccATGGCTGAAGAACCTACTACT |
| | 77 R | tccgccaccaccaaccactttgtacaagaaagctgggtaGAGAGGTTTGACATTGAACC |
| | 78 F | acaagtttgtacaaaaaagcaggctctccaaccaccATGTTTTTCTCCGGCGATC |
| | 78 R | tccgccaccaccaaccactttgtacaagaaagctgggtaGCTCAGATCAAAACCCGC |

··· continued ···

| Purpose | Name | Sequence (5'-3') |
|---|---|---|
| LCI assay | nLUC-CEPR2 F | GGTACCATGTCGAGAAGACCAGACCTC |
| | nLUC-CEPR2 R | GTCGACACTGTAATCTTTCCAGTTGTGTC |
| | NRT1.2-cLUC F | GGTACCATGGAAGTGGAAGAAGAGGTCTCA |
| | NRT1.2-cLUC R | GTCGACTTAGCTTCTTGAACCAGTTGATCTAT |
| Pull-down | CEPR2$^{KD}$ F | GGATCCCGTTACAGAGTTGTGAAGATACGT |
| | CEPR2$^{KD}$ R | GTCGACTACTGTAATCTTTCCAGTTGTGTC |
| | NRT1.2$^{Loop}$ F | ggatccATGTCTGGTTCTCGTTTCT |
| | NRT1.2$^{Loop}$ R | gaattcTTATTTCAGAACGATTTTAACG |
| | ubc32 F | ggatccATGGCGGATGAGAGGTATAATCGG |
| | ubc32 R | gtcgacTCTGTCATCGACTGGCTTCTGA |
| | ubc33 F | CGgatccATGGCAGAAAAAGCTTGTATAAAGCGT |
| | ubc33 R | gtcgacTAATGCTTCTTTCTTGTTCCTTCTCCT |
| | ubc34 F | CGgatccATGGCAGAAAAGGCCTGTATAAAACG |
| | ubc34 R | gtcgacTCCCTGTTTATTGTTCTTTTTCCTCTC |
| | ubc32 C93S F | TTTGAAATGCTCAAGCTAATCTTGGTGTTAG |
| | ubc32 C93S R | CTAACACCAAGATTAGCTTGAGCATTTCAAA |
| | ubc33 C87S F | GTCACTCATAGACAAGCTGATTTTCTTT |
| | ubc33 C87S R | AAAAGAAAATCAGCTTGTCTATGAGTGACT |
| | ubc34 C87S F | TGAAAATCACTCATAGATAAACTAATTTTCTTCTGT |
| | ubc34 C87S R | ACAGAAGAAAATTAGTTTATCTATGAGTGATTTTCA |
| *Xenopus* oocyte assay | CEPR2 F | tccccccgggATGGAAGTGGAAGAAGAGGTCTC |
| | CEPR2 R | cgggatccTTAGCTTCTTGAACCAGTTGATCTATACTT |
| | NRT1.2 F | gaagatctATGCGTTACAGAGTTGTGAAGATACGT |
| | NRT1.2 R | gactagtCTATACTGTAATCTTTCCAGTTGTGTCT |
| 35::*NRT1.2* | NRT1.2 F | ggatccATGGAAGTGGAAGAAGAGGTCTC |
| | NRT1.2 R | gtcgacTTAGCTTCTTGAACCAGTTGATCTATACT |
| | NRT1.2-GFP R | gtcgacGCTTCTTGAACCAGTTGATCTATACT |
| BiFC assay | YFP$^{N}$-CEPR2 F | CACCATGTCGAGAAGACCAGACC |
| | YFP$^{N}$-CEPR2 R | TACTGTAATCTTTCCAGTTGTGTC |
| | YFP$^{C}$-NRT1.2 F | CACCATGGAAGTGGAAGAAGAGGTCTCA |
| | YFP$^{C}$-NRT1.2 R | GCTTCTTGAACCAGTTGATCTAT |
| MbSUS Fragment deletion analysis | Cub-CEPR2 200 - 977 aa F | acaagtttgtacaaaaaagcaggctctccaaccaccATGTTAGCTCGCTCCAACTTGACC |
| | Cub-CEPR2 200 - 977 aa R | tccgccaccaccaaccactttgtacaagaaagctgggtaTACTGTAATCTTTCCAGTTGTGTC |
| | Cub-CEPR2 400 - 977 aa F | acaagtttgtacaaaaaagcaggctctccaaccaccATGCAAGTTGTTGAAGGATTCTGGTCT |
| | Cub-CEPR2 400 - 977 aa R | tccgccaccaccaaccactttgtacaagaaagctgggtaTACTGTAATCTTTCCAGTTGTGTC |
| | Cub-CEPR2 600 - 977 aa F | acaagtttgtacaaaaaagcaggctctccaaccaccATGTTGGGGTTGAGTATTTGCAGT |
| | Cub-CEPR2 600 - 977 aa R | tccgccaccaccaaccactttgtacaagaaagctgggtaTACTGTAATCTTTCCAGTTGTGTC |
| | Cub-CEPR2 1-717 aa F | acaagtttgtacaaaaaagcaggctctccaaccaccATGTCGAGAAGACCAGACCTC |
| | Cub-CEPR2 1-717 aa R | tccgccaccaccaaccactttgtacaagaaagctgggtaCTTCAACCACTTAACCGCCA |
| | Cub-CEPR2 1-802 aa F | acaagtttgtacaaaaaagcaggctctccaaccaccATGTCGAGAAGACCAGACCTC |
| | Cub-CEPR2 1-802 aa R | tccgccaccaccaaccactttgtacaagaaagctgggtaTGCGATTCCTTTTGCCGCT |
| | Cub-CEPR2 1-896 aa F | acaagtttgtacaaaaaagcaggctctccaaccaccATGTCGAGAAGACCAGACCTC |
| | Cub-CEPR2 1-896 aa R | tccgccaccaccaaccactttgtacaagaaagctgggtaCTCTCCAAACTCGTCTTCCA |
| | Cub-UBC32 F | acaagtttgtacaaaaaagcaggctctccaaccaccATGGCCGATGGTTTTGGTGAAC |
| | Cub-UBC32 R | tccgccaccaccaaccactttgtacaagaaagctgggtaAGACTGATCATCCATAAACC |
| | Cub-UBC33 F | acaagtttgtacaaaaaagcaggctctccaaccaccATGGCAGAAAAAGCTTGTATAAA |
| | Cub-UBC33 R | tccgccaccaccaaccactttgtacaagaaagctgggtaTCACAGCTGAAGCAAAGG |
| | Nub-NRT1.2$^{N}$ F | acaagtttgtacaaaaaagcaggctctccaaccaccATGGAAGTGGAAGAAGAGG |
| | Nub-NRT1.2$^{N}$ R | tccgccaccaccaaccactttgtacaagaaagctgggtaGAGAAAGATGAGAATAGAGACG |
| | Nub-NRT1.2$^{M}$ F | acaagtttgtacaaaaaagcaggctctccaaccaccATGGAAGACAACAAAGGATGGGA |
| | Nub-NRT1.2$^{M}$ R | tccgccaccaccaaccactttgtacaagaaagctgggtaTGGAGGTATCTTTAGGCTTC |
| | Nub-NRT1.2$^{C}$ F | acaagtttgtacaaaaaagcaggctctccaaccaccATGCTTCCGATATTTGCTTGC |
| | Nub-NRT1.2$^{C}$ R | tccgccaccaccaaccactttgtacaagaaagctgggtaGCTTCTTGAACCAGTTGATC |

⋯ continued ⋯

| Purpose | Name | Sequence (5'-3') |
| --- | --- | --- |
| Complementation | ProNRT1.2 F | aagcttatgtggtacgcttagtcaaattcaa |
| | ProNRT1.2 R | ggatccTCTCTCTCTTTCTTTCTCTCAAACTTT |
| | Overlap-NRT1.2$^{S292A/D}$-Fragment1 F | ggatccATGGAAGTGGAAGAAGAGGTCTC |
| | Overlap-NRT1.2$^{S292A/D}$-Fragment1 R | gcttcttcttgacgtggcttttccaattctccttgggcttcaaccttcttttttcccc |
| | Overlap-NRT1.2$^{S292A}$-Fragment2 F | ggggaaaaaagaagttgaagcccaaggagaattggaaaagccacgtcaagaagaagc |
| | Overlap-NRT1.2$^{S292A}$-Fragment2 R | gtcgacTTAGCTTCTTGAACCAGTTGATCTATACT |
| | Overlap-NRT1.2$^{S292D}$-Fragment1 F | ATTCTCCTTGGTCTTCAACTTCTTT |
| | Overlap-NRT1.2$^{S292D}$-Fragment2 R | AAAGAAGTTGAAGACCAAGGAGAATT |
